# Aquifer microbial communities differentially display metabolisms capable of secondary effects on uranium speciation across a former metal processing site

**DOI:** 10.64898/2026.06.01.729369

**Authors:** Catherine J. Pettinger, Allondra M. Woods, Raymond H. Johnson, Charles J. Paradis, Erica L.-W. Majumder

**Affiliations:** Department of Bacteriology, University of Wisconsin-Madison, Madison, WI 53706, USA; Biological Engineering Department, Utah State University, Logan, UT 84322, USA; RSI EnTech, LLC, Contractor to the U.S. Department of Energy Office of Legacy Management, Grand Junction, Colorado, USA; Department of Geosciences, University of Wisconsin at Milwaukee, Milwaukee, Wisconsin, USA

**Keywords:** Uranium Biogeochemistry, Microbiology Metabolism, Aquifer Communities, Subsurface Microbiology, Reduction-Oxidation Reactions

## Abstract

Groundwater contamination presents challenges across world, yet remediation solutions in variably oxidized regions are limited and many co-interactions between contaminant metals and microbial reactions occur. Here we present a genomic and metabolic study into the biogeochemistry of a uranium-contaminated surficial aquifer site in Riverton, WY. We identified unique communities that varied based on geochemistry, geography, and compartment, matching microbial subsurface studies. Cross-site metabolism tests showed communities had functional capabilities of nitrogen respiration, manganese reduction, iron reduction, and sulfide oxidization. No sites showed evidence of microbial U-bioreduction nor ammonium oxidation. Only former tailings area groundwater and ditch surface water sites nearest a retention pond, and a downgradient oxbow lake exhibited sulfate reduction metabolisms. This was contrary to our hypothesis of near-river downgradient groundwater sites having U and S reduction capability. Most communities which showed S reduction capacity exhibited Fe oxidation capacity. Modeling demonstrated U as calcium uranyl carbonates. Based on our metabolism tests and known mineral and microbial metabolism reduction potentials, this suggests U reduction could only be achieved via abiotic reaction with biogenic sulfide. Of eleven sites tested, it is possible in four. This has impact on future site-specific remediation plans and understanding of microbial reactions in variably reduced zones.

**Graphical Abstract:** 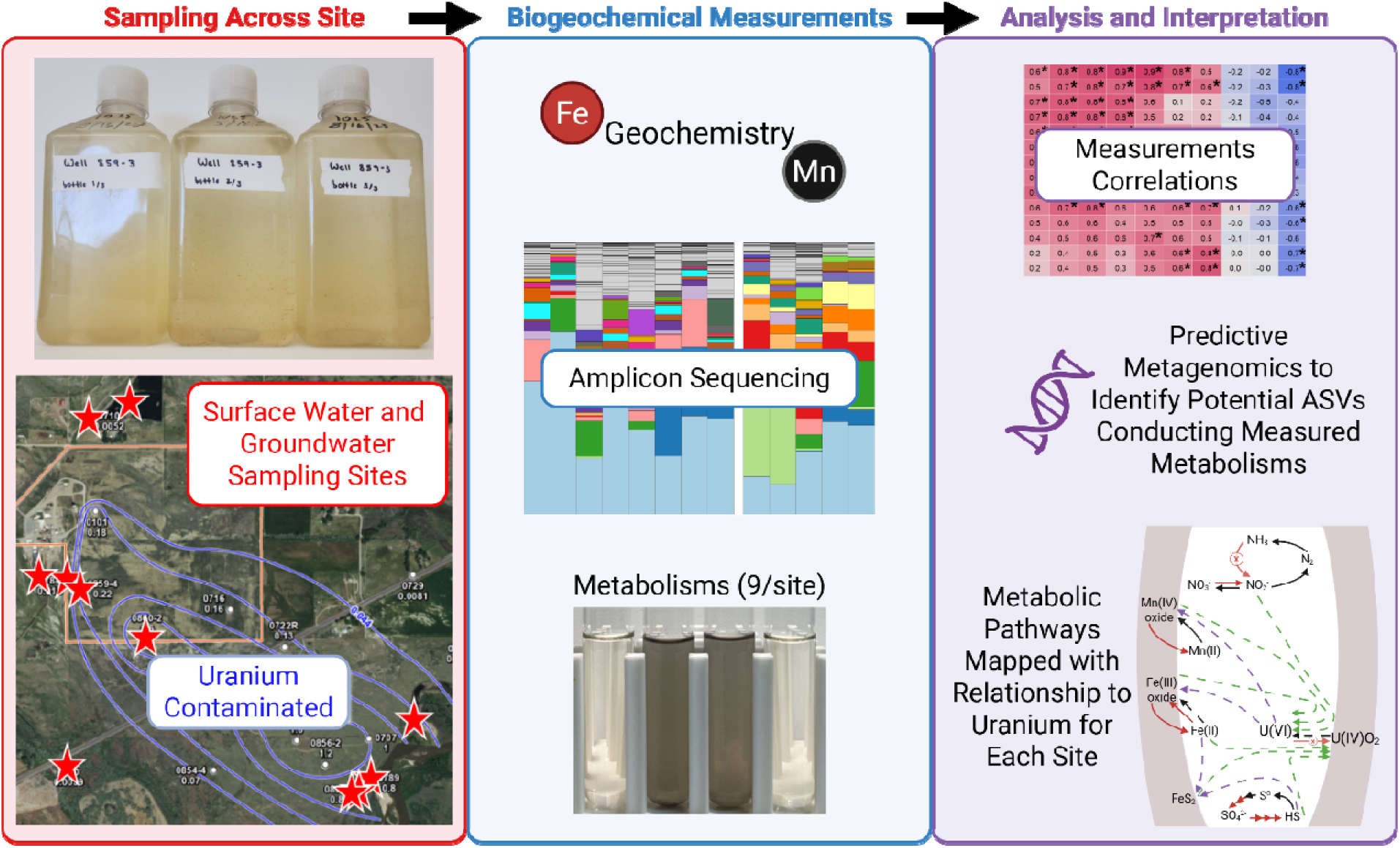

**Graphical Abstract Text:** We performed microbial membership and metabolism measurements across a uranium-contaminated site’s surface and ground waters, then performed analyses relating these metrics to geochemistry at the site. Findings showed variations in the membership, yet mainly similar functional capabilities. Metabolic differences were explained in relationship to uranium cycling and remediation implications.

## 1 Introduction

Groundwater contamination at uranium mill sites presents a biogeochemical challenge across the United States and the world. Access to freshwater is an issue recognized globally by the World Health Organization^1^, United Nations Environment Programme^2^, and more regionally in the Fifth National Climate Assessment which predicted water shortages to much of the United States, including the Western regions of the USA over the next 15 years^3^. The closure of World War II and Cold War era U mining, processing, and testing sites led to hazardous wastes at over 150 locations which persist and require continual monitoring^4^.

The Riverton WY Department of Energy (DOE) Legacy Management site is an example of an anthropogenic U contaminated freshwater aquifer, with former uranium and vanadium ore mill operations occurring from 1958 – 1963. During these operations, a tailings area of 72 acres of 4-foot depth with uranium, radium, and thorium was formed. 1.8 million cubic yards of waste was relocated in 1988 to the Gas Hills East Disposal Site with surface remediation performed in 1989^5^. However, the surficial aquifer was contaminated with U levels exceeding background levels of 0.044 mg/L^6^. Site levels are regularly monitored across the watershed and U is not flushing at the rate originally expected with detected concentrations varying before and after flooding events^6^. A map of the site is depicted as Figure 1. Water flows in the surficial aquifer southeast towards the Little Wind River^5^. The near-river region is referred to as the “SSMA”, or St. Stephen’s Mission Area. Once the water reaches the Little Wind River, U is diluted to below levels of human and ecological health concern and has low bioavailability^7^. An oxbow lake developed at the site in 1995^8^, which presents its own unique challenges for redox-sensitive metal transport dynamics^9–11^. Since the site has closed for uranium mill tailings, near the former tailings area (FTA) a sulfuric acid plant now operates with retention ponds of treated wastewater^12^ slightly upgradient from the main U plume.

**Figure 1.**
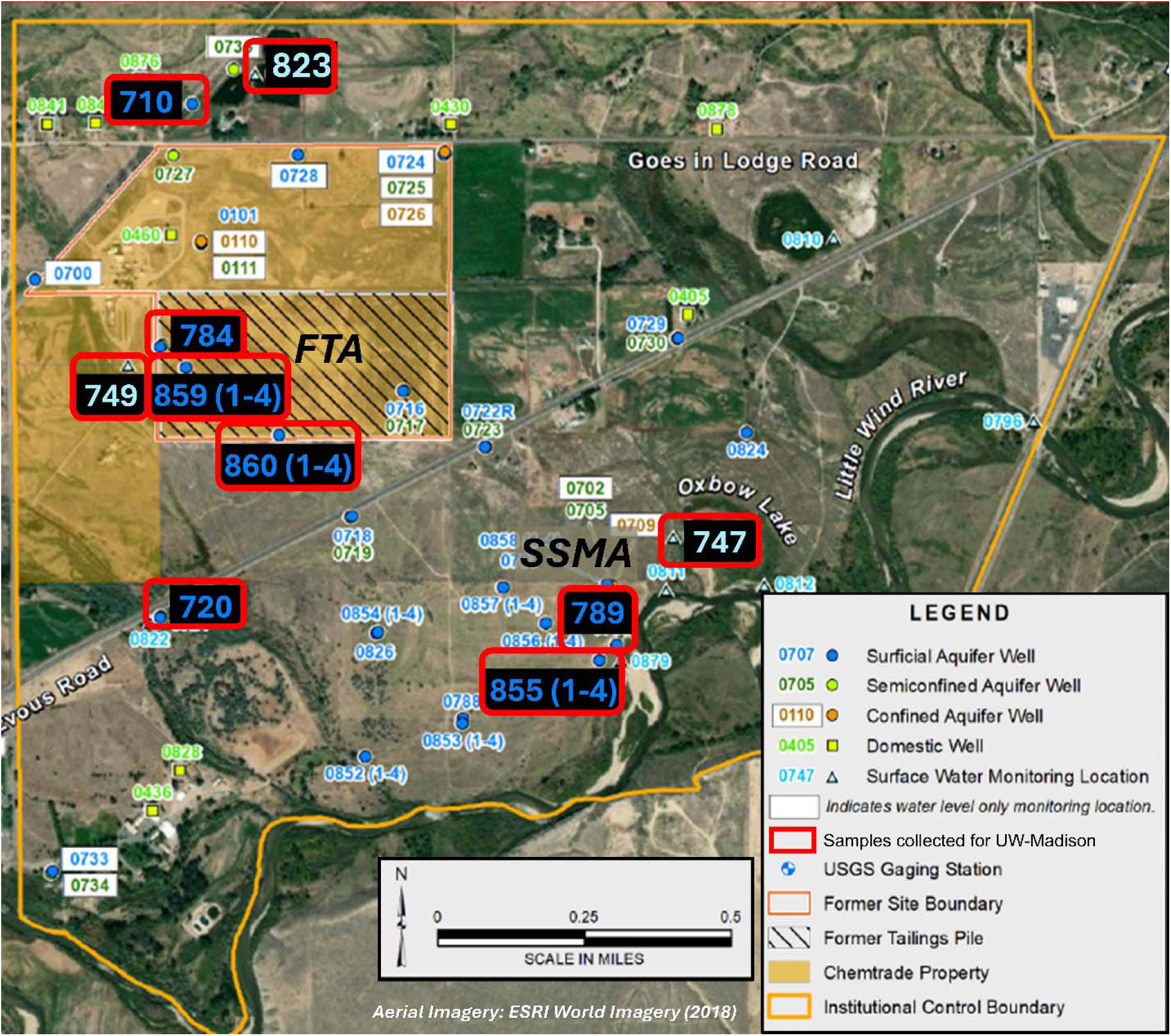
Sampling map of the Riverton site, adapted from 2023 Verification Monitoring Report^8^. The eleven sampling sites are indicated with a red box surrounding a blue number. Samples were collected from multi-channel wells 855, 859, and 860 and surficial aquifer wells 710, 720, 784, and 789. General groundwater flow is from the Former Tailings Area (FTA) area towards the Little Wind River, with site 710 described as “upgradient”, sites 784, 859, and 860 within the FTA, and sites 789 and 855 downgradient at the St. Stephens Mission Area (SSMA). Cross-gradient surficial aquifer site 720 and surface water sites 823 (pond), 749 (retention pond), and 747 (oxbow lake) were additionally sampled from.

Uranium mobility at the Riverton site have been recently explored. The Riverton site has a natural flushing plan based on estimated flushing within 100 years per CFR Code 40^13^, but recent models have suggested the flushing is not occurring at the rate expected^6^. Uranium remediation is often based on microbial reduction reactions to immobilize U^14–16^; generally U(VI) is more soluble compared to reduced U(IV)^17^. However, this strategy’s long-term efficacy is limited due to U(IV) reoxidation if reducing conditions are not maintained^18^. Instead of the traditional reductive immobilization scheme, oxidative enhanced flushing is utilized at this site. Oxidative enhanced flushing entails oxic bicarbonate-poor injections followed by oxic bicarbonate-rich injections^19^. The theory was to mobilize reduced U through oxidation, which has been shown in a previous study as more successful under carbonate facilitation^20^. Results from the trial showed that U, Fe, and Mn had differing results than advection-diffusion-dispersion tracer models alone could explain^19^. These transition metals are microbially influenced through reduction and oxidation metabolisms, and microbial reactions can exceed the abiotic reaction rate^21^ or bypass kinetic constraints^22^. However, it is unknown if microorganisms at this site are contributing to the unexpected U and transition metal responses to the oxic injections.

Riverton site microbial experiments have been limited to the SSMA region^23^ which we assert is not representative of the entire aquifer system. It is well-established at other metal-contaminated aquifer sites that microbial communities vary based on plume location, geochemistry, and compartment (sediment-bound versus planktonic)^24–26^, so we hypothesized that Riverton microbial communities would have similar physical drivers of their composition. We also hypothesized that functional tests would display regional variations. Specifically, we predicted microbial community members with known reductive metabolisms will be located closer to the river where there is a naturally reduced zone, and more oxidizing metabolisms in the FTA region. To test the hypotheses posed, we performed microbial community sequencing and nine metabolism stimulation tests on eleven resident Riverton water microbial communities. We then combined these results with predictive metagenomics utilizing PICRUST2^27^ and statistical analyses against geochemical data. Finally, we made predictive models of how these measured metabolisms interact with uranium. These experiments fill a site-specific knowledge gap; the Riverton site has not been tested for cross-site microbial membership nor metabolisms, and metal reduction-oxidation reactions, including microbial, are crucial to contaminant management strategy.

## 2 Materials and Methods

### 2.1 Materials and Reagents

Supplemental Table 2 provides the full list of reagents used and their manufacturer and catalog information.

### 2.2 Sampling

Three liters of unfiltered water were collected from multi-channel wells 855 (at depths “2” and “3”), 859 (depth “3”), and 860 (depth “3”), surficial aquifer wells 710, 720, 784, and 789, and surface water sites 823 (pond), 749 (retention pond), and 747 (oxbow lake) (Figure 1) during regular groundwater sampling of the Riverton WY site in August 2023 as described in US DOE Verification Monitoring Report, Riverton, WY, Processing Site^8^. Locations (elevation, latitude, longitude, depth) are recorded in Supplemental Table 1. Samples were collected into labeled sterile bottles alongside a shipping control with Luria Broth medium and shipped overnight in coolers with ice to the University of Wisconsin-Madison. Samples were stored at 4°C until use. No growth was noted in shipping control medium during sampling nor upon receipt of shipment.

### 2.3 Geochemical Measurements

Total groundwater iron (Fe) concentration was determined using inductively coupled plasma optical emission spectroscopy (ICP-OES, Agilent 5110 VDV) and an internal yttrium standard. Sample aliquots and calibrants were acidified with nitric acid (2%) prior to measurement. Fe concentration values were determined from Fe(II) chloride tetrahydrate calibrants (0.0900, 0.180, 0.890, 1.78, 3.11, and 4.44 ppm, R^2^ value of 1.00) in 2% nitric acid. All sample, calibrant, and blank measurements were determined based on the 238.20 nm wavelength.

Metrics for total alkalinity (as calcium carbonate), pH, specific conductance, sulfate, temperature, turbidity, and metal concentrations (Mn, Mo, U) were acquired from the Department of Energy (DOE) Geospatial Environmental Mapping System database^28^ (accessed June 12 2024) as reported in the US DOE 2023 Verification Monitoring report.^8^

### 2.4 Geochemical Modeling

PHREEQC version 3^29^ was utilized to model U and Mn chemical distributions at sites with noted differences in metabolism results by region. Sites 789, 855-2, 860-3, and 747 had geochemistry compiled from 2015 – 2019 from GEMS^28^, as those were the last dates that enough geochemical data surveyed for PHREEQC input requirements at these sites. Input data are listed in Supplemental Table 3. The database used was from Johnson et al., 2023 supplemental 6^30^ as this database has updated Ca and Mn carbonate complexes.

### 2.5 Stimulation of Microbial Reduction-Oxidation Metabolisms Tests

#### 2.5.1 Metabolism Test Solutions and Stocks Preparation

To test the Riverton groundwater microbial community across the site for the capability of 9 different energy metabolisms, raw groundwater was added to defined media and compared to sterilized (autoclaved) groundwater control additions to the same media. A base medium was designed by modifying the mineral, vitamin, and trace metal solutions described in Hurst et al.^31^ to approximate conditions in the Riverton aquifer and was used for all tests. Ion and metal concentrations were matched as feasible to averaged pre-injection levels across wells 1020 – 1034 in September 2020 and wells 859 and 860 [multi-depth] in June 2021 during field tracer testing at the site^19^. These sites and dates were chosen as they contain the most recent detailed metrics for the surficial aquifer including Cl, SO_4_, Ca, Mg, Na, K, I, Br, U, Mn, Mo, V, Sr, Se, Si, and Fe. The aquifer-defined medium (ADM) had 0.239 g/L ammonium sulfate, 0.100 g/L monobasic potassium phosphate, 0.500 g/L magnesium sulfate heptahydrate, 0.120 g/L calcium chloride dihydrate, 0.083 g/L sodium metasilicate pentahydrate, and 0.203 g/L yeast extract in Milli-Q (MQ) water. Trace metal solution was prepared by addition of 2.000 g/L nitrilotriacetic acid to MQ water, pH adjustment to 6.5, followed by addition of 1.000 g/L manganous sulfate monohydrate, 0.700 g/L iron(II) chloride tetrahydrate, 0.200 g/L cobalt(II) chloride hexahydrate, 0.200 g/L zinc sulfate heptahydrate, 0.020 g/L copper(II) chloride dihydrate, 0.020 g/L nickel chloride hexahydrate, 0.060 g/L sodium molybdate dihydrate, 0.020 g/L sodium selenate, 0.020 g/L sodium tungsten oxide dihydrate, and 0.005 g/L sodium orthovanadate. Vitamin solution consisted of 10 mg/L pyridoxine-HCl, 5 mg/L thiamine-HCl, 5 mg/L riboflavin, 5 mg/L calcium pantothenate, 5 mg/L thioctic acid, 5 mg/L *p*-aminobenzoic acid, 5 mg/L nicotinic acid, 5 mg/L vitamin B12, 5 mg/L mercaptoethanesulfonic acid, 2 mg/L biotin, and 2 mg/L folic acid in MQ water. Vitamin and trace metal solutions were adjusted to pH 7 and sterilized by filtration.

30.0 mL of vitamin solution and trace metal solution were dispensed to separate pre-autoclaved serum bottles and headspaces exchanged with a needle connected to 0.2 μm filter for ∼2 hours, swirling liquid throughout. Headspace-exchanged (“gassed”) solutions were used for all reduction metabolism tests. Non-gassed solutions were used for all oxidation metabolism tests. Biological atmosphere mixture for culture growth (Airgas Z03NI8522003062, 5% H_2_, 10% CO_2_, BAL% N_2_) was used for all gas exchanges described via a gas manifold.

50 mL serum bottles of 1.00 M sodium bicarbonate were bubbled for 20 minutes with a needle inserted into solution, headspaces gassed for additional hour, and autoclaved. 0.500 M sodium nitrate, 0.100 M Fe(III) citrate, and 0.500 M Mn(IV) oxide, and 0.050 M uranyl acetate stocks were prepared by bubbling MQ water for 30 minutes, then weighed solute added. Gas-exchange continued for an additional 15 minutes. Nitrate, Fe(III), and Mn(IV) stocks were autoclaved. Uranyl acetate was filter sterilized (0.2 μm Nalgene RapidFlow) and headspace-exchanged with a 0.2 μm filtered needle.

#### 2.5.2 Metabolism-Specific Stimulation Additions

Metabolism tests performed are summarized in Table 1, with test-specific reactant and carbon additions alongside endpoint method described.

**Table 1.** Metabolism-Specific Stimulation Additions and Endpoint Methodology.

| Metabolism Test | Abbr. | Reactant Added | Carbon Sources | Endpoint |
| --- | --- | --- | --- | --- |
| Ammonium Oxidation | AO | 18.1 mM ammonium sulfate | 0.080 g/L peptone, 0.24 mM sodium acetate | Quantitative: NH<sub>4</sub><sup>+</sup> depletion by IC |
| Mn(II) Oxidation | MnO | 0.12 mM manganous sulfate | 0.080 g/L peptone, 0.24 mM sodium acetate | Quantitative: aqueous Mn concentration by ICP-MS/MS |
| Sulfide Oxidation | SO | 38 mM sodium thiosulfate | 0.080 g/L peptone, 12 mM sodium acetate | Quantitative: SO<sub>4</sub><sup>2-</sup> formation by IC |
| Fe(II) Oxidation | FeO | Bottom medium: 50mM iron(II) chloride tetrahydrate | Bottom medium: 1.00 mM sodium acetate | Qualitative: Fe(III) banding pattern, gradient culture |
| Nitrate Respiration | NR | 10 mM sodium nitrate | 20 mM sodium DL-lactate, 10 mM sodium acetate | Quantitative: NO<sub>3</sub><sup>-</sup> depletion by IC |
| Mn(IV) Reduction | MnR | 20 mM manganese oxide | 7.1 mM sodium DL-lactate, 1.6 mM sodium acetate | Quantitative: Mn(IV) depletion by benzidine assay |
| Fe(III) Reduction | FeR | 8 mM iron(III) citrate | 2.5 mM sodium DL-lactate, 0.88 mM sodium acetate | Quantitative: Fe(II) production by ferrozine assay |
| U(VI) Reduction | UR | 2 mM uranyl acetate | 2.5 mM sodium DL-lactate, 0.88 mM sodium acetate | Quantitative: U(VI) depletion by spectrophotometry assay |
| Dissimilatory Sulfate Reduction | DSR | 30.0 mM sodium sulfate | 60.0 mM lactate, 1.00 g/L yeast extract | Quantitative: $\text{SO}_4^{2-}$ depletion by IC, Qualitative: Black Precipitates |

Ammonia oxidation (AO), and sulfide oxidation (SO) metabolism tests were prepared by addition of reactant and carbon sources to ADM per Table 1. For SO metabolism test, sodium thiosulfate was used as substrate instead of sulfide based on Luo et al.^32^ These media were autoclaved, and post-autoclave 10.0 mL/L vitamin solution, 10.0 mL/L trace metal solution, and 20.0 mL/L 1.00 M sodium bicarbonate were added. pH was adjusted to 7.2 ± 0.2. 10.0 mL media dispensed to sterilized borosilicate culture tubes (18 × 150 mm).

Manganese(II) oxidation (MnO) tests were prepared by addition of carbon sources per Table 1 to ADM. This MnO medium was autoclaved. Post-autoclave additions were filter-sterilized manganous sulfate per Table 1 alongside other post-autoclave additions, pH adjustment, and aliquoting to culture tubes as described for AO and SO media.

Fe(II) oxidation (FeO) was tested in gradient Fe(II)-oxygen cultures based on procedures described in Sobolev & Roden^33^, Emerson & Merrill Floyd^34^, and Yli-Hmeminki et al.^35^. A bottom layer medium was prepared with 1.00 mM sodium acetate, and 2% w/v agar in ADM. A top layer medium was prepared with ADM. The pH was adjusted for both top and bottom media to 7.2 ± 0.1. Media were autoclaved, caps on media were tightened upon autoclave door opening, and further steps were performed in an anaerobic Coy chamber (Coy Laboratory Products, Grass Lake, Michigan, USA) with mixed gas (Airgas X03NI73C30014T7, 7% H_2_, 20% CO_2_, BAL% N_2_). 2.0 mL/100.0 mL 1.00 M sodium bicarbonate and 10.0 mL/100.0 mL 0.500 M FeCl_2_ were added to the bottom layer medium. 1.0 mL bottom layer medium was dispensed to sterilized borosilicate culture tubes (13 x 100 mm) and allowed to solidify under anaerobic atmosphere for ∼1 hour. Top layer medium was cooled to 30°C, and 10.0 mL/L vitamin solution, 10.0 mL/L trace metals, and 20.0 mL/L sodium bicarbonate were added to the top layer medium. 4.0 mL of top layer medium was added to the solidified bottom layer medium. These solidified overnight in the anaerobic chamber, then were removed to aerobic laboratory settings to form the O_2_ gradient culture conditions. 20.00 μL inoculum was injected as a pipette was pulled through the top media. This was performed for: each site in duplicate (replicates R1 and R2), each site’s autoclaved groundwater control, and a MQ water control.

Nitrate respiration (NR), Mn(IV) reduction (MnR), Fe(III) reduction (FeR), and U(VI) reduction (UR) metabolisms were tested. 9.60 mL of ADM base medium was added to Hungate-style tubes and condition-specific volumes of sodium acetate and sodium DL-lactate were added (Table 1). Media were gas-exchanged (Airgas Z03NI8522003062, 5% H_2_, 10% CO_2_, BAL% N_2_) 15 - 20 minutes utilizing a gas manifold equipped with long needles reaching the bottoms of tubes, then crimp-sealed with butyl stoppers and autoclaved. Post-autoclave, 0.10 mL vitamin solution, 0.10 mL trace metal solution, and 0.20 mL 1 M sodium bicarbonate solution were added to each tube. Reactants were then added per Table 1 with stocks as prepared in 2.4.1. pH was adjusted to 6.9 - 7.0. Nitrate usage was determined based on Cardenas et al.^36^, Westrop et al.^37^ and Fries et al.^38^. Fe(III) citrate usage by Cardenas et al.^36^ and Cordoso et al.^39^ Uranyl acetate usage by Jeon et al.^40^

The dissimilatory sulfate reduction (DSR) metabolism test used MOYLS4 medium^41^, prepared with 8.00 mM magnesium dichloride hexahydrate, 20.0 mM ammonium chloride, 0.600 mM calcium chloride dihydrate, 2.00 mM dipotassium phosphate, 2.00 mM monosodium phosphate, 1.00 g/L yeast extract, 15.0 mL/L 2.00 M Tris-HCl (pH 7.4), 0.060 mM FeCl_2_, 0.120 mM EDTA, 60.0 mM lactate, and 30.0 mM sodium sulfate anhydrous in MQ water. The pH was adjusted to 7.2. 9.80 mL was dispensed to Hungate-style tubes, gas-exchanged for 15 – 20 minutes as described for other reduction tests, and autoclaved. 0.10 mL of vitamin and trace metal solutions were then added to each DSR tube.

#### 2.5.3 Inoculation of Metabolism Tests

All metabolism tests, except for FeO that was inoculated as described in 2.4.2, were inoculated with 1.0 mL of raw groundwater to a labelled Replicate 1 (R1) and Replicate 2 (R2) tube for each site for each media type upon opening of sampling bottles in laboratory. 1.0 mL of autoclaved/sterilized (S) groundwater for each site for each media type was also prepared. 1.0 mL of autoclaved MQ water was prepared for each media. All inoculations were performed next to a flame. All metabolism tests were incubated at room temperature in dark and were inverted or lightly vortexed three times per week.

#### 2.5.4 Endpoints of Metabolism Tests

All qualitative endpoints were assessed at 6 weeks. All quantitative endpoints were performed after 15 weeks. Images were taken throughout tests with 2 – 3 images/metabolism test/site/week on a regular schedule.

Qualitative metabolism endpoints included DSR (also determined quantitatively as described next) and FeO. DSR metabolisms were determined by formation of black Fe(II) sulfide precipitates^36,41^. FeO metabolisms were determined by formation of an Fe-oxide ring in the gradient culture as described by previous literature^33–35^.

NR, AO, SO, and DSR metabolisms were determined quantitively by changes in anion or cation concentration as listed in Table 1 by ion chromatography (IC). IC procedure used a Dionex 4 mm ICS-2100 and ICS-1100 with mobile phases of 15 mM methanesulfonic acid and 3.5 mM sodium carbonate/1 mM sodium bicarbonate, a 25 μL loop size, AS14/CS12A columns, and ASRS 4 mm/CSRS_4mm suppressors. Sulfate concentrations were determined from peak area (μS × min) at ∼9.4 minutes against a calibration curve (4.00 × 10^1^, 2.00 × 10^2^, 3.60 × 10^2^, 5.20 × 10^2^, 6.80 × 10^2^ ppm calibrants, R^2^ value of 0.998). Ammonium concentrations were determined from peak area (μS × min) at 3.75 minutes against a calibration curve (6.00, 30.0, 54.0, 78.0, and 102 ppm calibrants, R^2^ value of 0.999). Nitrate concentrations were determined from peak area (μS × min) at 6.43 minutes against a calibration curve (6.00, 30.0, 54.0, 78.0, and 102 ppm calibrants, R^2^ value of 0.996). As needed, 10× or 20× dilutions for samples were performed.

Mn(IV) reduction was assessed quantitatively by the benzidine assay adapted from Burnes et al.^42^ Briefly, 50.0 µL sample was added to 1.00 mL 2.00 mM benzidine hydrochloride solution in 10% acetic acid, 200.0 μL of this added to a plate reader, and the absorbance was read at 415 nm over 20 minutes at 30 second intervals. Maximum absorbance values were compared to calibrants prepared from Mn(IV) oxide (2.00, 5.00, 10.0, 20.0, 30.0 mM calibrants, R^2^ value 0.996) and oxidation state selectiveness by benzidine was verified by a Mn(II) chloride tetrahydrate control.

MnO metabolism was to be assessed quantitively initially with the benzidine assay^42^. The benzidine assay was unable to accurately measure the lower Mn(IV) concentrations needed for this study, and total aqueous Mn levels were instead determined by inductively coupled plasma tandem mass spectrometry (ICP-MS/MS). Since measurement was taken after filtering, we assumed that the sparingly soluble Mn(IV) oxides were removed and only Mn(II) remained in the solution that was measured. ICP-MS/MS (Agilent 1260 Infinity II) was performed with 0.22 μm syringe-filtered sample that was diluted (20×) and acidified (2% HNO_3_) against prepared manganous sulfate standards (10.0, 100., 200., 300., 400., 500. ppb, R^2^ value 1.00). An ICP-MS/MS internal yttrium standard was used. Each sample was measured in triplicate by the instrument.

FeR metabolism was assessed quantitatively by the ferrozine assay^39^ for formation of Fe(II). Briefly, 100.0 μL sample was added to 1.00 mL 0.500 M HCl. 200.0 μL of this was added to 1.30 mL ferrozine solution (1.00 g/L ferrozine in 50.0 mM HEPES). After 10 minutes, the solution was 0.2 μm syringe-filtered, and absorbance was read at 562 nm. Each sample was measured in duplicate, compared against Fe(II) standards (1.00 mM, 2.00 mM, 4.00 mM, 6.00 mM, 8.00 mM, 10.00 mM, R^2^ 0.991) prepared with a FeCl_2_/EDTA stock, and values averaged.

UR metabolism was assessed by U(VI) spectrophotometric determination.^43,44^ Briefly, 10.0 μL sample was added to 100. μL complexing solution (25 g/L DCTA, 5 g/L sodium fluoride, 65 g/L sulfosalicylic acid, pH 7.85), 100. μL Tris buffer (pH 7.85), 500. μL ethanol, 100. μL PADAP (100 g 2-(5-Bromo-2-pyridylazo)-5-(diethylamino)phenol per 100. mL ethanol), and 350. μL MilliQ water. The mixture was incubated for 40 minutes, and absorbance was measured at 578 nm. Each sample was measured in duplicate, compared against U(VI) standards (0.20 mM, 0.30 mM, 0.50 mM, 1.00 mM, 2.00 mM, 3.00 mM, R^2^ value 1.00) prepared from uranyl acetate, and the values were averaged.

Percent change for metabolism tests were determined quantitatively by disappearance of the reactant (AO, NR, MnO, MnR, DSR, and UR). This was calculated as:

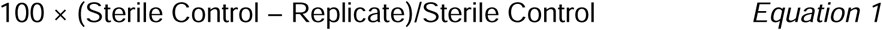

The sterile control used in this calculation was the site-specific sterile control as measured alongside all replicates. Replicate percent change values were then averaged for each site.

Percent change for metabolism tests determined by appearance of product was calculated as:

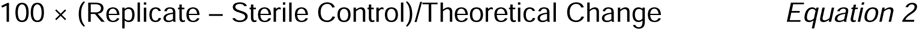

This was performed for FeR using a theoretical change of 8 mM, based on an expected maximum expected yield change of 8 mM Fe(III) (added as reactant) converting to Fe(II) product. Theoretical change value for SO was 76 mM based on reactant addition of 38 mM sodium thiosulfate (Na_2_S_2_O_3_) converting to SO_4_. Replicate percent changes were then averaged for each site.

Percent change for binary FeO test was performed by averaging the binary result (0 or 1) and multiplying that by 100. This resulted in values of 0, 50, or 100 for FeO percent change options, where 0 corresponds to no Fe oxidation detected, 50 to one replicate with FeO detected, and 100 to both replicates with FeO detected.

### 2.6 DNA Extraction and Amplicon Community Sequencing

Approximately 1 liter of water from each site was filtered through 0.1 μm a PES membrane filters (ThermoScientific Nalgene RapidFlow Lot 1376832). Filter membrane was removed with an autoclaved razor blades and tweezers, stored in a sterile tube, and frozen at - 80°C. Filters for sites 747 and 823 were split in further extraction and sequencing steps.

DNA was extracted using Qiagen DNeasy PowerWater Kit instructions. DNA concentrations were read via a Qubit 3 Fluorometer (Invitrogen by ThermoFisher Scientific) using broad range solutions (1× dsDNA BR assay kit, ThermoFisher).

Amplicon library preparation with PCR-amplification targeting the V4 region of the 16S rRNA gene and sequencing on an Illumina Miseq was performed according to our established methods^45^. Briefly, primers used were dual-indexed designed using high-fidelity polymerase^46^, Pfx. A negative control and positive control (Pfx) were added during each sequencing run. Gel electrophoresis was performed verifying amplified PCR products. Amplified products were normalized to 20 μL using a SequalPrep Normalization kit (Life Technologies) and aliquots of 5 μL from normalized samples were pooled into the final library. Final concentrations verified using KAPA library quantification kit (Kapa Biosystems) and a Qubit 2.0 Fluorometer (Invitrogen). Then, the final library was diluted to 20 nM with HT1 buffer and PhiX control v3 (20% vol/vol) and 600 μL loaded onto a MiSeq v2 (500 cycles) reagent cartridge (Illumina). Sequences were uploaded to Illumina Sequence Hub.

### 2.7 Microbial Community Analyses

Sequencing files were accessed using BaseSpace Sequence Hub Downloader (Illumina) and processed for bioinformatics via Quantitative Insights into Microbial Ecology (QIIME) 2 (version 2024.5)^47^. Data was uploaded to Qiime2 using Casava 1.8 paired-end demultiplexed format (via Qiime import tools). DADA2^48^ denoise-paired action was used to trim and truncate sequences (values chosen based on demux file interactive quality plot for below 8 or past the 30% quality control threshold), denoise, and quality filter (with chimera consensus method). Amplicon sequence variants (ASVs) were aligned using MAFFT^49^ and FastTree^50^ (both via Qiime-phylogeny). Taxonomic assignments were made with a Naïve Bayes Classifier, trained by Sklearn 1.4.2 on 05/30/2024 using Silva 138 99% operational taxonomic unit full-length sequences (provided from Qiime2)^51–53^.

Qiime2-processed files were uploaded in R (version 4.4.0)^54^ using qza_to_phyloseq via qiime2R. These were then processed and compositional barplots visualized using package microViz^55^. Alpha diversity was calculated using phyloseq^56^ function estimate_richness for Shannon’s diversity index. Beta diversity was performed with vegan^57^ function metaMDS for Bray-Curtis dissimilarity index for two dimensions (k = 2) and visualized with ggplot as part of tidyverse^58^ with metadata of water type and location descriptors added.

Microbial community function predictions were performed using PICRUST2 (version 2.6.2)^27^. Key outputs from this were functional abundances of enzyme classifications and MetaCyc^59^ pathway coverages.

### 2.8 Correlation Analyses

Correspondence analysis between geochemistry values in Supplemental Table 3 and microbial abundance in Supplemental Table 5 were performed in R using the vegan^57^ package. Amplicon sequence variants (ASVs) were pre-filtered with the criteria that they were present in at least three samples including site replicates, and the constrained correspondence analysis function was used with default settings e.g. double-standardization for Chi-square inertia as described in vegan package documentation (version 2.6-6.1)^57^. The CCA plot was visualized using the ggplot package^58^. Spearman correlations were calculated and visualized for ASVs against geochemical values (Table 2), and ASVs against Supplemental Table 7 percent change values, using the microViz package^55^. ASVs were prefiltered such that replicate samples were removed and ASVs were required to be present in three sites for correlation analysis.

**Table 2.** Geochemical values for sampling sites. Iron values determined from ICP-OES and other metrics compiled from 2023 Verification Monitoring Report^8^ and GEMS Legacy Management database^28^.

| Site Code | Location Descriptor | Total Alkalinity as CaCO<sub>3</sub>, ppm | Mn, ppm | Mo, ppm | pH | Specific Conductance, µmhos/cm | Sulfate, ppm | Temp °C | Turbidity, NTU | U, ppm | Fe, ppm |
| --- | --- | --- | --- | --- | --- | --- | --- | --- | --- | --- | --- |
| *Groundwaters* |  |  |  |  |  |  |  |  |  |  |  |
| 710 | Upgradient | 188 | 0.0184 | 0.00311 | 7.27 | 719 | 146 | 11.57 | 0.29 | 0.00523 | 0 |
| 720 | Crossgradient | 226 | 0.002 | 0.00201 | 7.21 | 568 | 78.2 | 13.5 | 0.53 | 0.00386 | 0 |
| 784 | FTA | 370 | 0.79 | 0.167 | 7.26 | 4161 | 1830 | 16.96 | 1.62 | 0.0137 | 0.02 |
| 859-3 | FTA | 305 | 1.03 | 0.0965 | 6.84 | 6047 | 3280 | 15.3 | 3.47 | 0.145 | 3.19 |
| 860-3 | FTA | 248 | 1.44 | 0.26 | 6.82 | 3741 | 1930 | 15.34 | 1.11 | 0.981 | 1.64 |
| 855-2 | SSMA | 435 | 0.441 | 0.309 | 7.08 | 9240 | 5030 | 15.11 | 1.86 | 0.825 | 2.28 |
| 855-3 | SSMA | 388 | 0.839 | 0.329 | 7.03 | 8336 | 4420 | 15.26 | 2.92 | 0.755 | 0.09 |
| 789 | SSMA | 365 | 0.0822 | 0.491 | 7.01 | 7927 | 4375 | 12.73 | 0.65 | 0.8655 | 0.02 |
| *Surface Waters* |  |  |  |  |  |  |  |  |  |  |  |
| 823 | Upgradient | 95 | 0.0533 | 0.000673 | 8.78 | 2757 | 966 | 27.28 | 3.82 | 0.00298 | 0.01 |
| 749 | Retention Pond | 179 | 0.0172 | 0.00455 | 7.17 | 954 | 225 | 20.59 | 6.4 | 0.000137 | 0.91 |
| 747 | Oxbow Lake | 323 | 0.698 | 0.0185 | 9.33 | 1349 | 321 | 18.38 | 13.1 | 0.101 | 0.09 |

## 3 Results

### 3.1 Riverton Geochemistry

Geochemistry values^28^ are recorded in Table 2. U was elevated above site MCL of 0.044 mg/L at FTA sites 859-3 and 860-3, all groundwater SSMA sites, and oxbow lake site 747. Aqueous Mn and Mo were detectable in all sites, however metals Mn and U were highest at FTA site 860-3 whereas Mo was highest in SSMA sites. Fe in the groundwater samples varied and was undetectable in 710 and 720, whereas within the plume concentrations were detected. Sulfate levels ranged across the site with upgradient and cross-gradient sites at 146 and 78.2 mg/L, respectively, and plume groundwater concentrations elevated ranging from 1830 – 5030 mg/L. pH was roughly circumneutral within groundwater samples, ranging from 6.82 – 7.27 and slightly elevated in upgradient surface water and the oxbow lake. Turbidity was highest at oxbow lake site, 747, and lowest at upgradient pond site, 710. These results show the Riverton as pH circumneutral with high sulfate concentrations and varying metal concentrations based on the U plume.

PHREEQC^29^ was utilized to determine U speciation for sites SSMA sites 789 and 855-2, FTA sites 859-3 and 860-3, and oxbow lake site 747 based on site archived geochemistry data (Supplemental Table 3). Sites were chosen for analysis based on metabolism test results discussed later. Distributions of U species showed oxidation state 6 calcium uranyl carbonates primarily at all sites (Supplemental Table 4) and manganese as oxidation state 2.

### 3.2 Riverton Microbial Community

1076 total unique ASVs were identified in the water from the 11 Riverton well sites sampled (Supplemental Table 5). For the alpha diversity indices observed amplicon sequence variants (ASV), Chao1, and Abundance-based Coverage Estimator (ACE), values per site varied from 69 – 151 (Supplemental Table 6) and total counts per location ranged from 2050 – 7378 (Supplemental Table 5). FTA sites had higher diversity than SSMA sites based on richness and evenness indices Shannon and Simpson, despite richness indices Chao1, observed ASVs, and ACE indices for SSMA sites being within the range of FTA site values (Supplemental Table 5). Beta diversity metric Bray-Curtis dissimilarity was used to compare microbial communities across the Riverton samples (Figure 2). Communities ordinated across the x-axis based on surface versus groundwater locations. Samples within subgroups FTA and SSMA were more similar to each other than other locations, although most samples were not strongly clustered together. Replicates from surface water sites 823 and 747 were relatively similar to each other rather than the groundwater sites. Notably, retention pond site 749 was furthest from other samples, suggesting higher dissimilarity in the planktonic community of the retention pond from nearby groundwater region FTA. Results show alpha diversity differences between site regions and beta diversity differences by water type (groundwater versus surface water) and site region.

**Figure 2.**
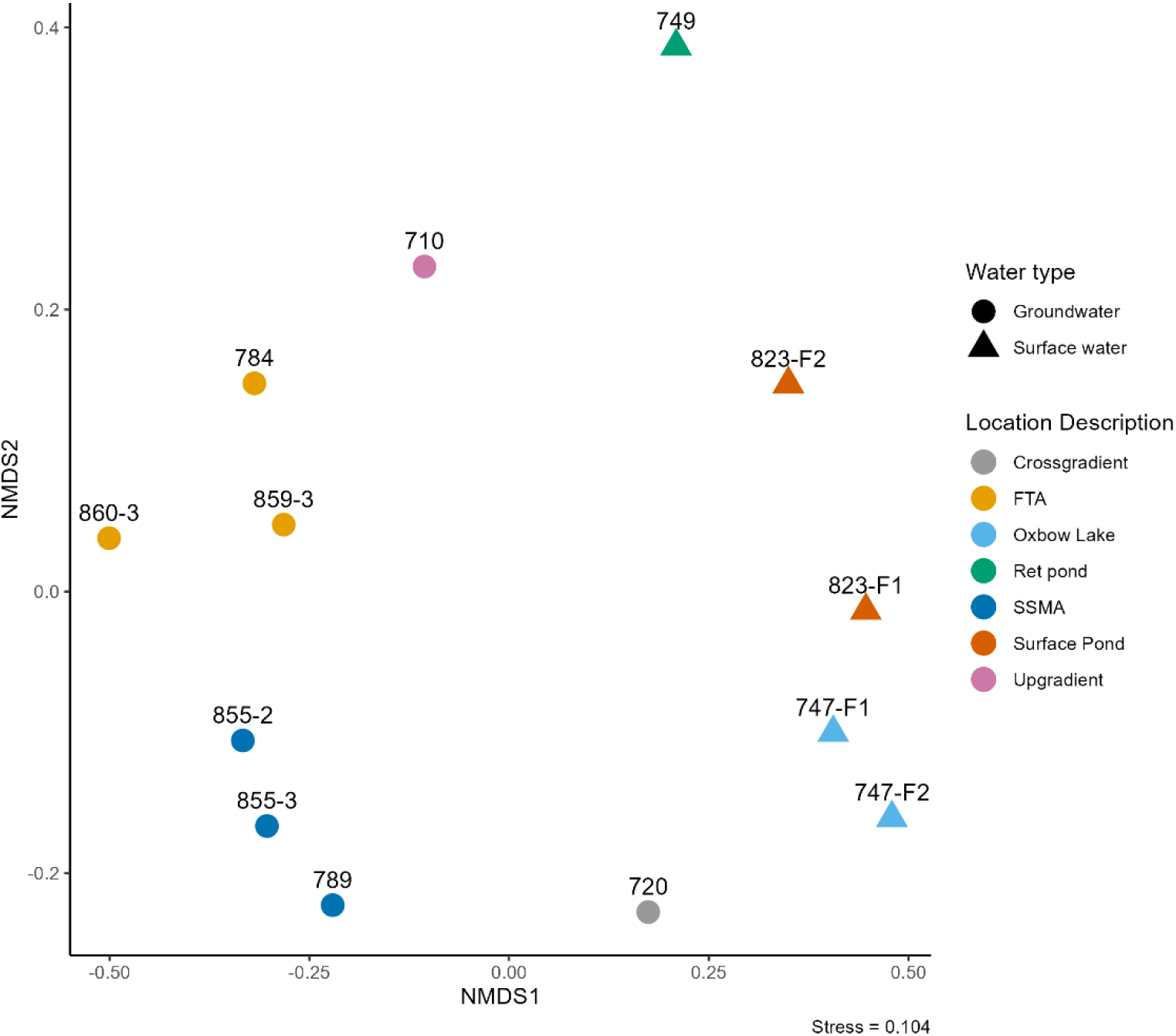
NMDS plot of Bray-Curtis Dissimilarity Beta diversity results comparing the water microbial communities across Riverton site. Descriptors used for site locations are indicated by color, and groundwater versus surface water by shape.

Microbial community membership was compared between samples (Figure 3, Supplemental Table 5) and some taxonomic orders were found throughout all sampling locations. At the order level, ASVs identified as Burkholderiales were present in all groundwater and surface water sites. This was mainly family Comamonadaceae. No other families of Burkholderiales were found throughout all samples. Order Pseudomonadales was identified in all sites, although minimally present in site 747. Families of this order included Cellvibrionaceae, Moraxellaceae, Pseudomonadaceae among others, but no family within the order was identified throughout. Order Flavobacteriales was found in all regions (FTA, SSMA, upgradient, crossgradient, surface waters) but was missing from SSMA sites 855-2 and 855-3. Order Rhodobacterales was found in all regions as well but was missing from FTA site 784. Sequencing results showed similarities across locations for some taxonomy orders but few lower classifications within these orders were the same between sites.

**Figure 3.**
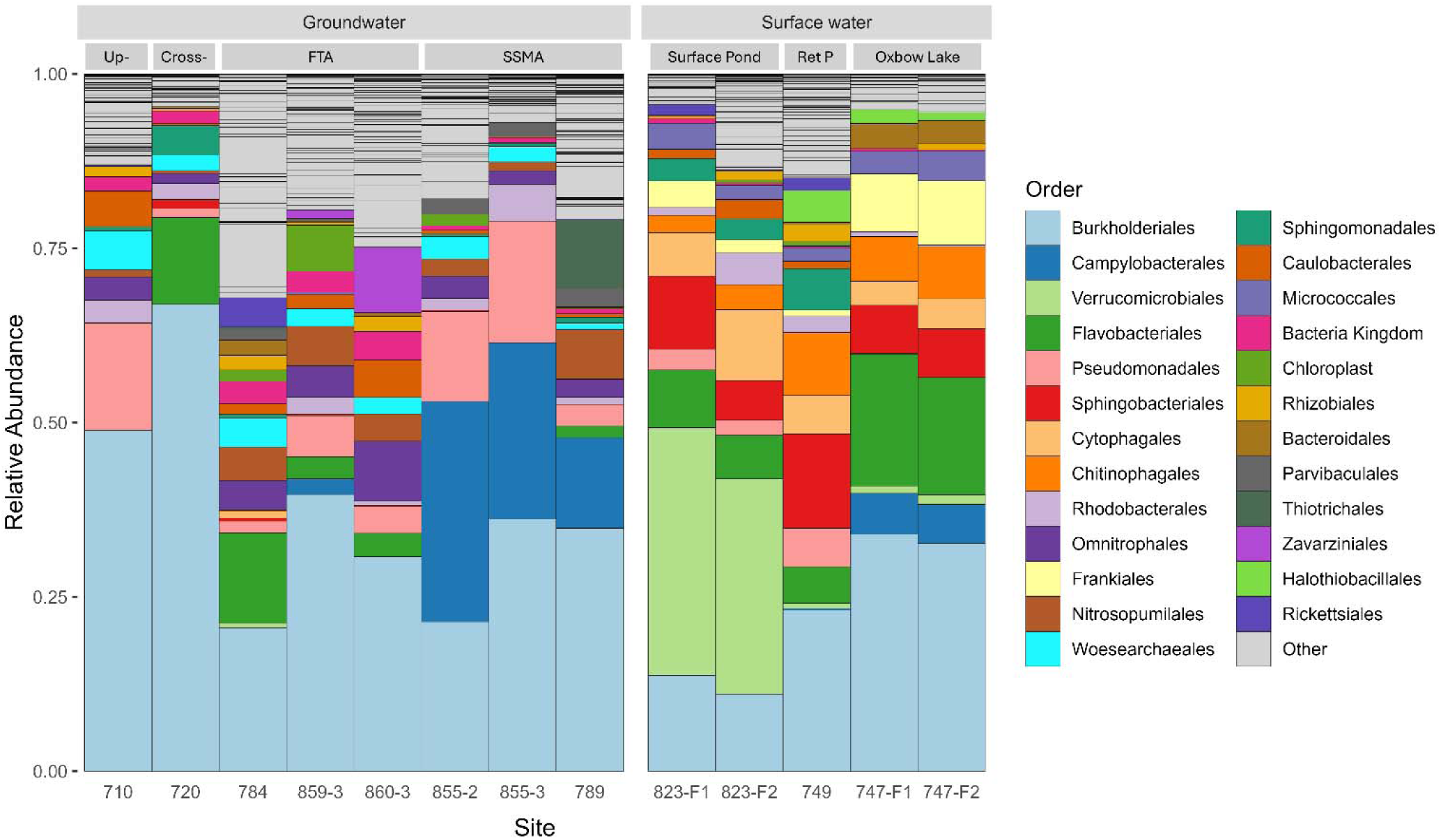
Relative abundance of the microbial community members from across the Riverton site. The top 25 orders identified by 16S amplicon sequencing are displayed. Samples are sorted by groundwater (surficial aquifer) and surface water samples. Groundwater samples are further sorted by their relationship to the U plume, with upgradient, cross-gradient, former tailings area (FTA), and St. Stephen’s Mission Area (SSMA) zones labelled. Surface water sites include two replicates of the upgradient surface pond and downgradient oxbow lake site and one replicate of the retention pond near the FTA zone.

Surface waters had differing microbial communities compared to groundwaters. Surface waters had higher relative abundances of Sphingobacteriales (surface waters 7 – 13%, groundwaters 0 – 1.3%), Cytophagales (surface waters 3 – 10%, groundwaters 0 – 1%), Chitinophagales (surface waters 3 – 9%, groundwaters 0 – 0.2%), Frankiales (surface waters 1 – 9%, groundwaters 0%) and Micrococcales (surface waters 2 – 4%, groundwaters 0 – 0.3%). Sphingomonadales order was present in upgradient pond (3%) and retention pond (6%), although this order was not present in the oxbow lake and minimally in groundwaters (<1%). Groundwaters presented higher relative abundances than surface waters orders Woesearchaeales (1 – 6% groundwaters, 0% surface waters), Omnitrophales (1 – 9% groundwaters, 0 – 0.1% surface waters), and Nitrosopumilales (0.5 – 7% groundwaters, 0% surface waters).

Regionally the microbial community varied. In the SSMA, ASVs identified as order Campylobacterales are present in both groundwater (sites 855-2, 855-3, and 789) and the surface water of the oxbow lake (747-F1 and 747-F2), whereas this order was minimally present in the FTA. The relative abundance between groundwater and surface water of Campylobacterales varied though, with groundwater SSMA sites at 13 – 32% relative abundance compared to the oxbow lake at 6%. FTA and upgradient groundwater site 710 had higher values of Caulobacterales than SSMA sites (1.5 – 5% compared to SSMA at 0.1 – 0.5%) and Rhizobiales (0.4 – 2%, SSMA 0 – 0.2%). Desulfitobacteriales was present in FTA sites 784 and 859-3 and oxbow lake site 749 but was not present in the other samples.

Sites also had specific orders associated with them. For example, Thiotrichales (identified at genera level as *Thiothrix*) was present at 10% in site 789 but less than 0.3% in all other sites. Halothiobacillales (identified at genera level as *Thiovirga*) is present at 5% in site 749, minimally present (1 – 2%) in site 747, and not present in any other sites. Rickettsiales was present at 4% in site 784, minimally present in sites 823 and 749 (1.5% and 1.8%, respectively), present at <1% in sites 710 and 789, and 0% at other sites. Lachnospirales was present at 9% in site 784, present at <1% sites 859-3 and 789, and not present in any other sites.

Thalassabaculales (identified at genera level as *Thalassobaculum*) was present at 4% in site 860-3, 0.3% in 855-2, and undetected in other sites. Many other orders were detected with minimal presence across the site or were dispersed at various locations, e.g. CHAB-XI-27, Legionellales, Reyranellales, Pirellulales, Desulfobulbales, etc. 16S amplicon sequencing of the microbial community show variability based on water type and geographical location, with sites closer together having some underlying similarities but each with distinct microbial membership.

### 3.3 Correlation of Geochemistry and Microbial Community Composition

To further analyze how the planktonic microbial composition may vary based on site geochemistry, two correlation analyses were performed. Canonical Correspondence Analysis (CCA) results (Figure 4A) show that the abundance of multiple taxa share a set of geochemical drivers including Fe, Mn, sulfate, U, Mo, and conductivity. These ASV include members from families Terasakiellaceae, Gallionellaceae, Rhodocyclaceae, Nitrospiraceae, Nitrosopumilaceae, Omnitrophaceae, and others. Other microbial families, e.g. Sporichthyaceae, Cellvibrionaceae, Alteromanadaceae, and Bacteroides, appear to be driven by turbidity, pH, and temperature parameters.

**Figure 4.**
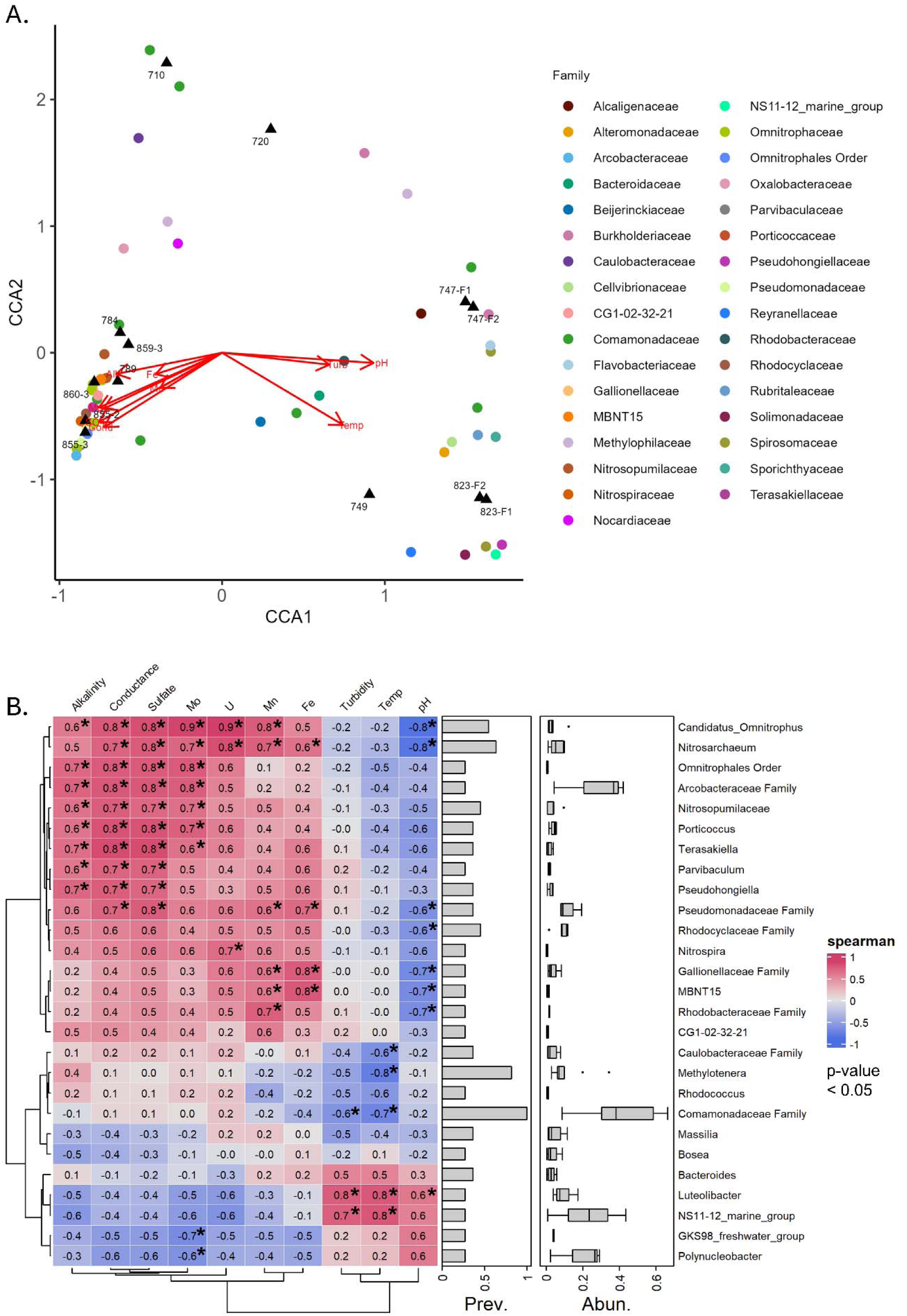
Canonical correspondence analysis (CCA) (A) and Spearman correlations (B) of the most prevalent amplicon sequence variants (ASVs) and geochemistry. A) CCA plot. ASVs, shown as circles, were filtered based on presence in at least 3 sites and each ASV is colored based on its family-level designation. Sites are oriented on the CCA plot based on their geochemistry and are represented as black triangles. Red arrows depict geochemical drivers. **B)** Spearmen correlation plot. Value of each correlation is shown and colored (positive correlations as red, negative correlations as blue). Prevalence and abundance of each ASV is shown, and ASV are described on the genera level or the lowest identifiable level. *p*-values from the Spearman correlation with a value <0.05 are marked with an asterisk.

Multiple microbial members were found to have significant positive Spearman correlations with alkalinity, conductance, sulfate, Mo, U, Mn, and Fe (Figure 4B). These same microbial members are negatively correlated with pH, with genera *Candidatus Omnitrophus, Nitrosarchaeum, Nitrospira,* and *MBNT15* showing significance. ASVs without genera resolution identified as families Psuedomonadaceae, Rhodocyclaceae, Gallionellaceae, and Rhodobacteraceae similarly demonstrate significant positive correlations with metal abundance and negative correlation to pH. Taxonomic abundance within the Riverton site was related to geochemistry with multiple cross-correlations, as shown in both two-variable (geochemical/taxon) results from the Spearman matrix and the multivariate CCA.

### 3.4 Microbial Metabolism Tests

Metabolism tests were performed to determine potential for Riverton microbial communities to carry out metabolisms and uncover any variability in functional potential. Ammonium oxidation (AO) occurs through sequential oxidation steps of ammonium to nitrite then to nitrate (Figure 5A, reactions 5 – 7). Aerobic AO metabolism was tested by measuring the disappearance of ammonium from the medium compared to sterile controls (Supplemental Figure 1A). The ammonium concentration differences between sterile controls and site-specific replicates were small, with all values ranging from 22 – 24 mM ammonium (Figure 5B) and percent change values within +/- 5% (Supplemental Table 7), suggesting that ammonium oxidation did not occur in these tests or rates of ammonium oxidation matched those of reactions producing ammonium by other members of the microbial community e.g. N_2_ fixation, DNRA, or mineralization. AO tests were performed aerobically, so it is unlikely that DNRA occurred in the presence of oxygen as a terminal electron acceptor or that anammox occurred.

**Figure 5.**
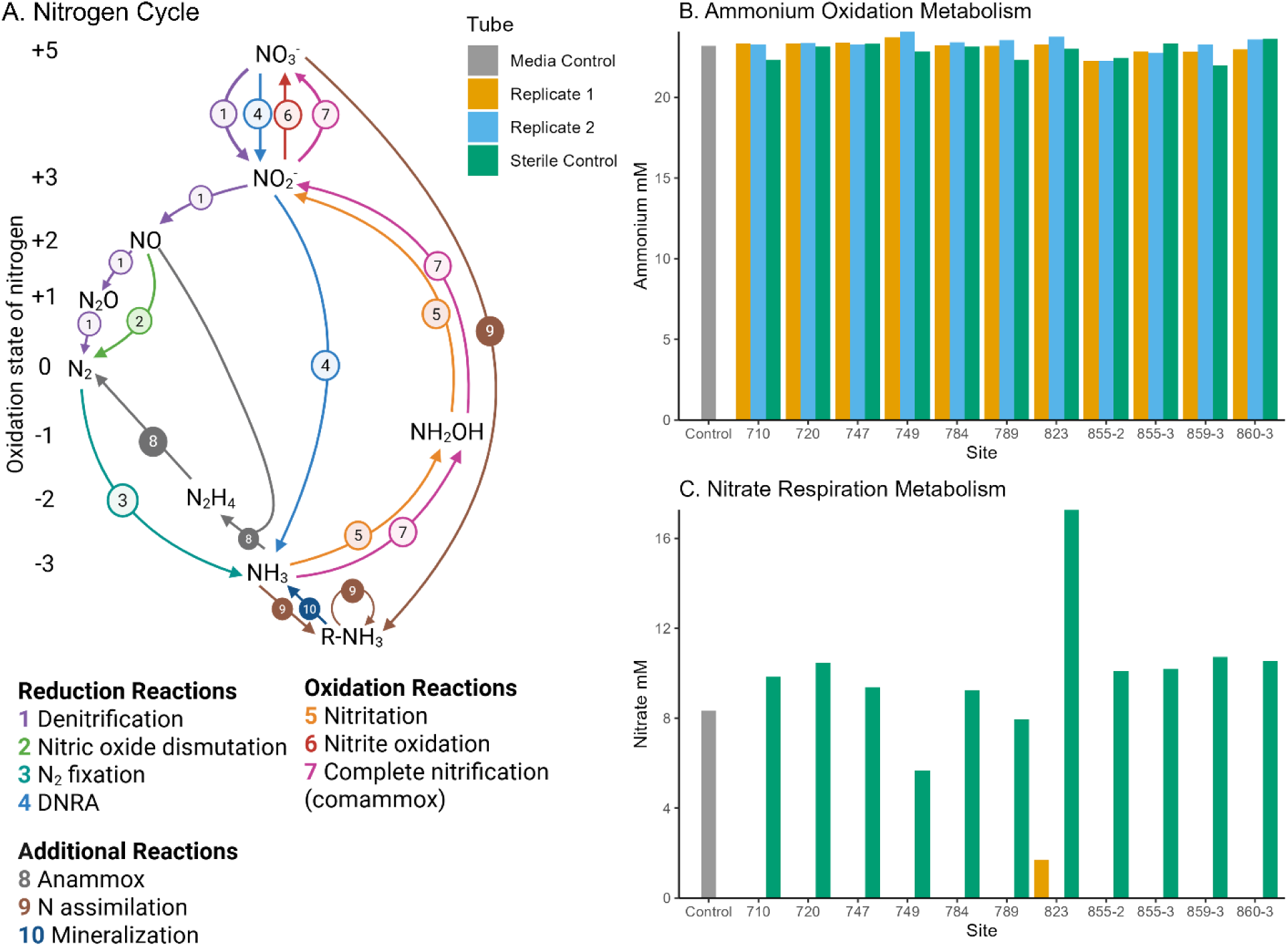
Nitrogen cycle metabolism results. A) Nitrogen cycle as adapted from Zhang et al. 2020^60,61^ and Kuypers et al. 2018^62^, displaying microbial transformation reactions of major environmentally-relevant nitrogen oxidation states (+5, +3, +2, +1, 0, -1, -2, -3) and generalized assimilated N (R-NH_3_) which can exist as various oxidation states. B) AO metabolism test results with ammonium concentration in mM. C) NR metabolism test results with nitrate concentration in mM. Bars not shown are levels of nitrate undetectable by ion chromatography.

Nitrate respiration (NR) includes denitrification (Figure 5A reaction 1) and dissimilatory nitrate reduction to ammonium (DNRA, Figure 5A reaction 4). NR was tested quantitatively by nitrate disappearance from the medium (Supplemental Figure 1B). Results (Figure 5C) show complete nitrate depletion at every site except site 823 which had a 95% change of nitrate (Supplemental Table 7). Notably, nitrate concentrations varied across site sterile controls, with surface water site 823 at 17 mM being almost double the concentration of other locations. The results demonstrate cross-site microbial potential of nitrate reduction when stimulated across the Riverton site samples.

Fe(III) reduction (FeR) metabolism was tested via measurement of Fe(II) production (Supplemental Figure 1F). Fe(II) was produced in most sites except for one replicate from cross-gradient groundwater site 720, upgradient groundwater site 710, and the autoclaved controls (Figure 6A). This suggests that most sites within the groundwater U plume are capable of FeR when stimulated.

**Figure 6.**
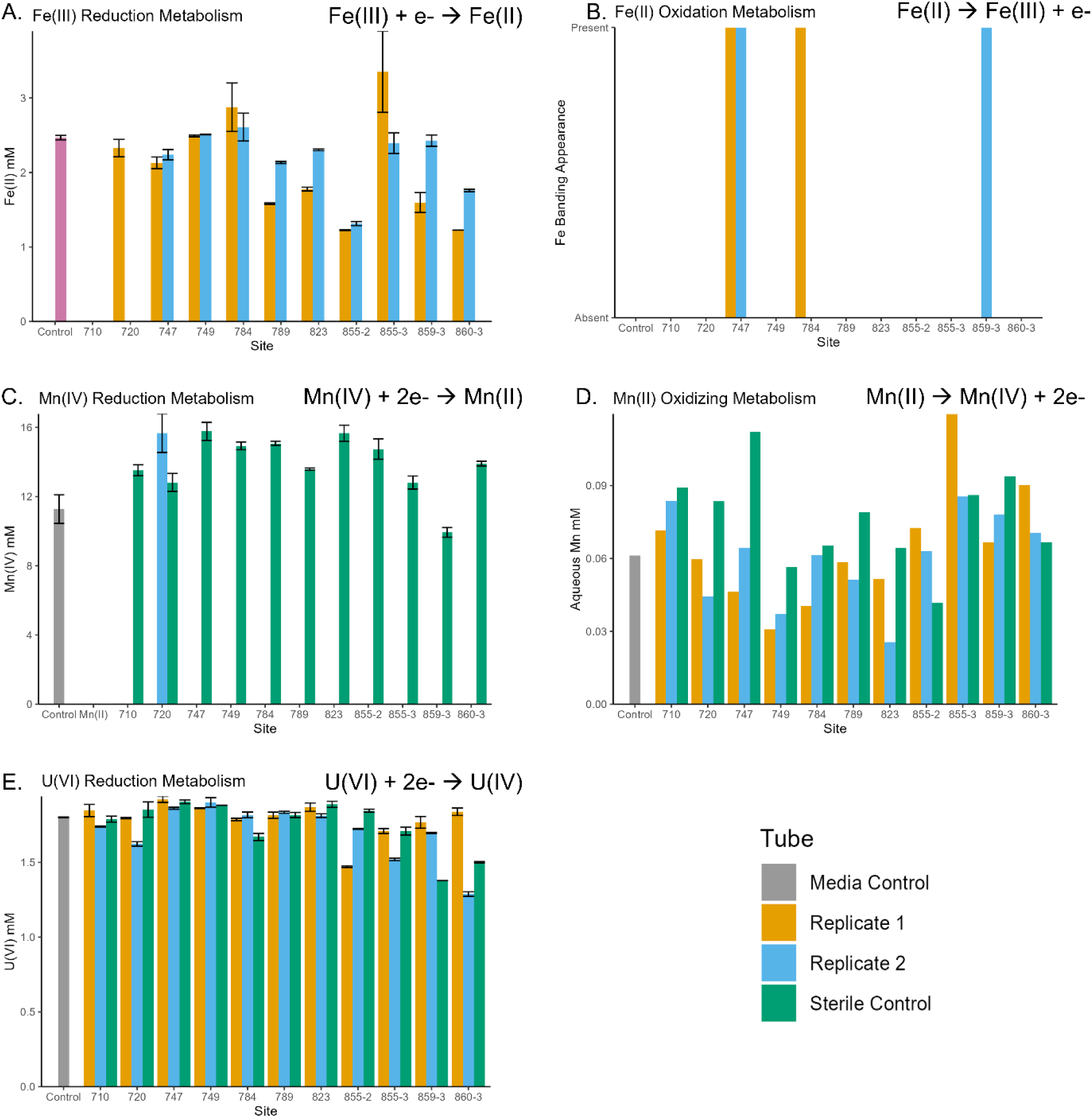
Metal metabolism results for Fe, Mn, and U. A) Fe reduction metabolism measured Fe(II) concentration. B) Fe oxidation metabolism are binary presence/absence (1/0) data from banding appearance in Fe(II) – O_2_ gradient cultures. C) Mn(IV) reduction based on Mn(IV) concentrations. D) Mn(II) oxidation was assessed based on aqueous Mn concentration. E) U(VI) reduction tests based on U(VI) concentrations. Error bars present for Fe(III), Mn(IV), and U(VI) represent technical replicates of spectrophotometric assay results. Bars not plotted were values less than assay calibrant range.

Fe oxidation (FeO) metabolism was tested through a gradient culture (Supplemental Figure 1E). Results were qualitative based on formation of reddish-brown Fe band compared to controls with gradient as described in previous literature^33–35^. Fe bands were observed in both replicates of oxbow lake site 747 and one replicate of FTA sites 784 and 859-3 (Figure 6B & Supplemental Figure 1E). This suggests that the Riverton microbial community across the site, unlike FeR, had a limited capacity for FeO in the conditions tested.

Mn reduction (MnR) metabolism was measured with Mn(IV) disappearance (Supplemental Figure 1D). MnR results (Figure 6C) show complete Mn(IV) disappearance in all sites and replicates. The one exception was a replicate in the cross-gradient of U plume groundwater site 720. The sterile controls showed little to no disappearance of Mn(IV), as expected (Figure 6C, green bars).

Mn oxidation (MnO) metabolism was tested through disappearance of Mn(II) (Supplemental Figure 1C). MnO results (Figure 6D) had variability between replicates with some replicates exceeding aqueous Mn concentrations of the sterile controls. The amount of Mn measured in the sterile controls across all samples also varied more than would be expected given concentration in Riverton groundwater used from each site. This indicated that MnO may have occurred but potentially obscured by active cycling between Mn(II) and Mn(IV); as such, MnO metabolism specifically was unable to be assessed in the methodology utilized herein.

Uranium reduction (UR) metabolism was tested through the disappearance of U(VI) (Supplemental Figure 1G). Some replicates show a small decrease relative to sterile controls (Figure 6E), but only site 855-2 had both replicates positive for UR, whereas in 859-3 both replicates had higher concentrations of U(VI) than the control at the end point suggesting possible oxidation. Therefore, microbial UR metabolism was not overtly demonstrated by the planktonic microbial community across the Riverton site.

Sulfide oxidation (SO) was tested quantitatively by production of sulfate compared to sterile controls (Supplemental Figure 1H). Thiosulfate was added in excess, which can be oxidized to sulfate and elemental sulfur or terathionate (Figure 7A, reaction 3). Elemental sulfur (S^0^) can then be oxidized by multiple pathways to sulfite and sulfate as detailed in Figure 7A. In all sulfide oxidation metabolism replicates, sulfate production occurred and sulfate levels exceeded those of controls (Figure 7B). These results suggest that sulfide oxidation by the microbial community can be stimulated across the site.

**Figure 7.**
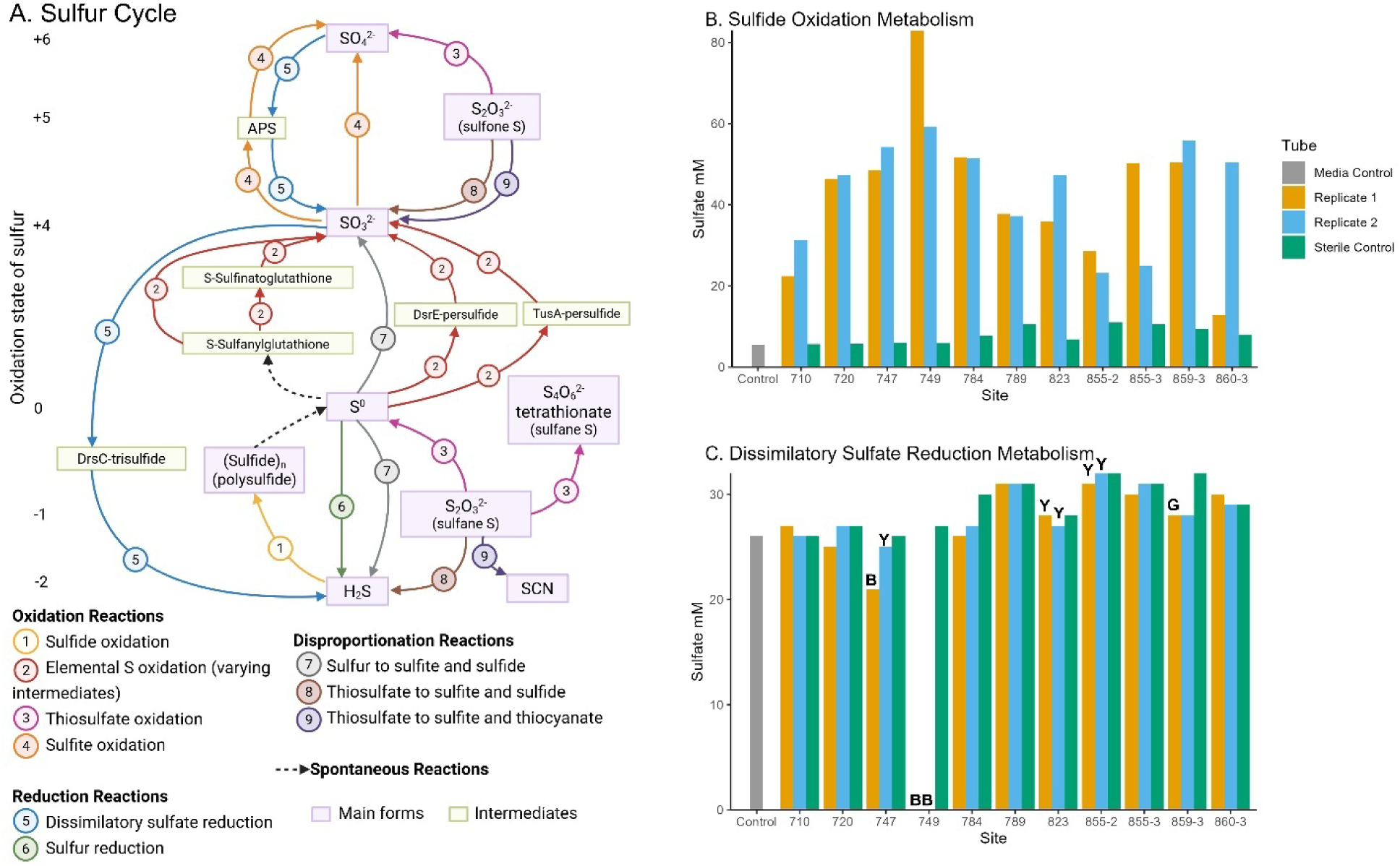
Sulfur cycle metabolism results. A) S cycle, simplified from Zhou 2025^63^ and displays microbial oxidation, reduction, and disproportionation reactions of major environmentally-relevant sulfur oxidation states (+6, +5, +4, 0, -1, -2). B) Sulfide oxidation (SO) metabolism test results with sulfate concentration in mM. C) Dissimilatory sulfate reduction (DSR) metabolism test results with sulfate concentration in mM. Bars not shown are levels of sulfate undetectable by ion chromatography. Qualitative results are indicated for DSR metabolism with “B” indicating black precipitates, “Y” yellow precipitates, and “G” gray coloring of media.

Dissimilatory sulfate reduction (DSR) was tested quantitatively by a decrease in sulfate concentration (Supplemental Figure 1I). As noted in Figure 7A, sulfate is reduced by DSR metabolism to sulfite then to hydrogen sulfide. There were notable drops in sulfate concentration in samples 747, 749, 784, and 859-3 (Figure 7C). These had average sulfate percent-change values of 12%, 100%, 12% and 12%, respectively (Supplemental Table 7). DSR was also assessed qualitatively by the appearance of black Fe(II) sulfide precipitates. There was distinct blackening of the medium by the 6-week timepoint of 747 Replicate 1 and 749 both replicates, and there was slight graying of 859-3 replicate 1 (Supplemental Figure 1I). Yellow precipitates formed in 747 replicate 2, 823 replicate 2, and 855-2 both replicates (Supplemental Figure 1I). Sites 710, 720, 784, and 860-3 did not have DSR metabolism noted through qualitative nor quantitative metrics.

Metabolism results showed the microbial community across the site to be capable of nitrate reduction (NR), Mn reduction (MnR), and sulfide oxidation (SO). Most communities were also capable of Fe reduction (FeR). No sites were shown as capable of direct, complete, U(VI) reduction (UR) or ammonium oxidation (AO). Differing sites were capable of dissimilatory sulfate reduction (DSR) and Fe oxidation (FeO).

### 3.5 Correlational Test of Microbial Community Membership and Metabolism Endpoints

To test if measured amounts of different microbial metabolisms were related to the abundance of microbial community members, a spearman correlation was performed (Figure 8). *Nitrospira* had a significant positive correlation with ammonium oxidation, although percent change values for AO metabolism were relatively low, ∼5% (Supplemental Table 7).

**Figure 8.**
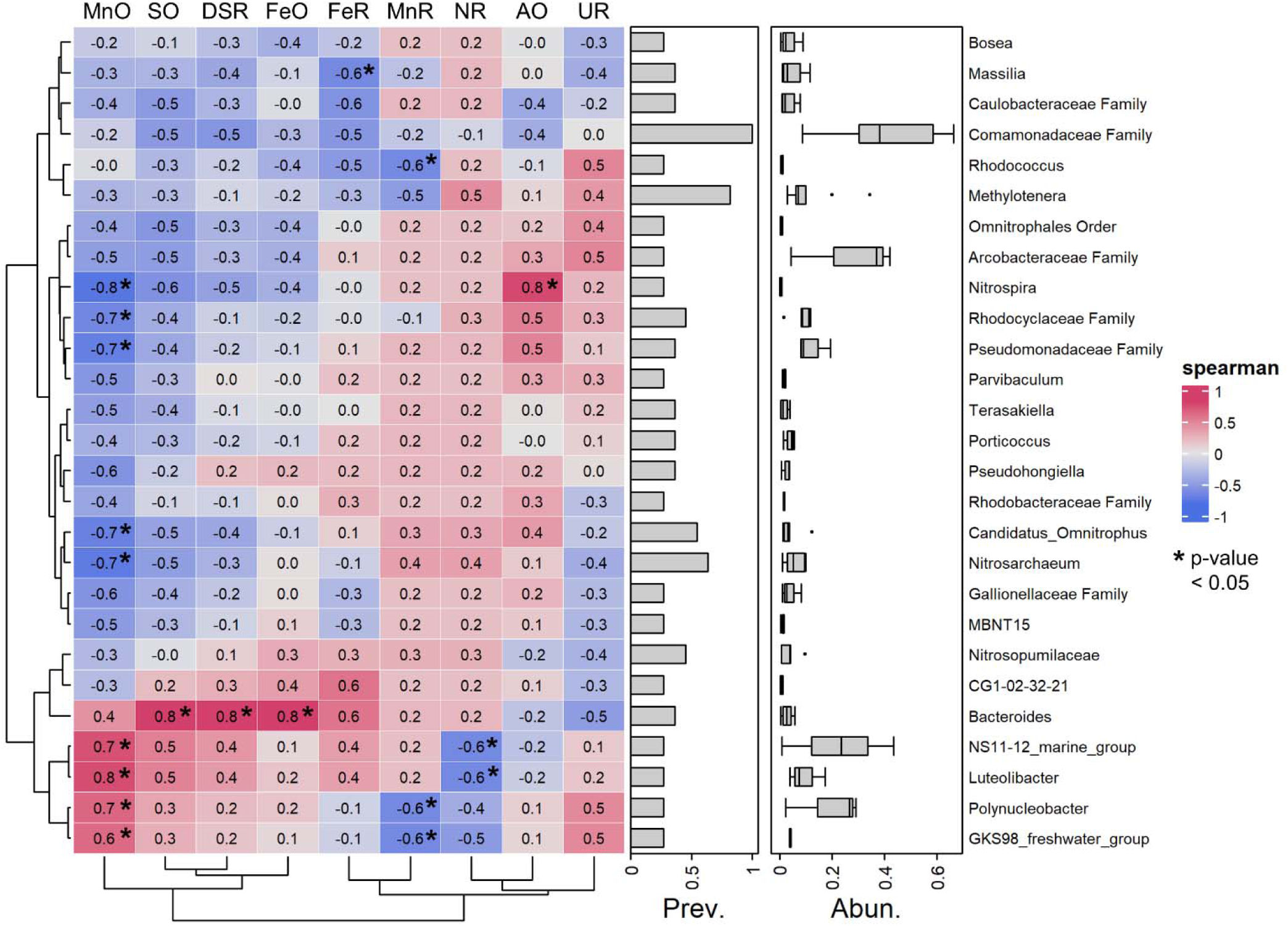
Spearman correlation between reduction-oxidation metabolism test percent change values and amplicon sequence variant (ASV) abundance. Samples 747 and 823 second replicates were removed and ASVs were filtered to be present in at least 3 samples for consideration. The value of each correlation is shown and colored (red-blue) based on value. Prevalence and abundance of each ASV is shown, and ASV are described on the genera level or the lowest identifiable level. *p*-values from the spearman correlation with a value <0.05 are marked with an asterisk.

*Bacteroides*, an obligate anaerobe, had significant positive correlations with SO, DSR, and FeO metabolisms. Although MnO had the most either significant positive or negative correlations with nine bacterial ASVs, the percent change (Supplemental Table 7) values of MnO metabolism were inconsistent between replicates and against controls. There were some significant negative correlations noted, which indicated that the presence of these microbes may be inhibitory to that metabolism, e.g. *Massilia*, *Polynucleobacter* and *GKS98 freshwater group* to Fe reduction, *Rhodococcus* to Mn reduction, and *NS11-12 marine group* and *Luteolibacter* to nitrate reduction. Otherwise, there were slight but insignificant correlations (*p* > 0.05) with other ASVs and metabolisms.

### 3.6 Predictive Metabolisms of Community Members

Microbial members had metabolic reactions predicted using PICRUST2^27^ to the nearest member metagenomic dataset available (Supplemental Table 8). These were used to predict metabolic pathway coverage per site sampled (Supplemental Table 9). 63% of pathways predicted were present in all sites, and most pathways present at high values were present at all sites (e.g. PWY0-1586 [peptidoglycan maturation], PWY66-389 [phytol degradation], NONOXIPENT-PWY [pentose phosphate pathway (non-oxidative branch) I], and PWY-1042 [glycolysis IV]).

E.C. predictions showed differing metabolic capabilities per ASV (Supplemental Table 8). For example, GKS98_freshwater_group was predicted to have E.C. 1.16.3.3, part of manganese oxidation II pathway. Other ASVs identified as positively correlated with MnO metabolism (Figure 8) were not predicted by PICRUST2 to have this metabolism. No ASVs were predicted with E.C. 1.16.9.1 [iron-oxidizing protein Cyc2]. Only 18 ASVs were predicted as having either of the AO pathway enzymes E.C. 1.14.99.39 or 1.7.2.6. 773 ASVs, however, were predicted as having either or both DSR pathway reactions E.C. 2.7.7.4 or E.C. 1.8.99.2, corresponding to the first and second arrow, respectively, of Figure 7A reaction 5. This included *Bacteroides*, although *Bacteroides* was not predicted as having E.C. 1.8.5.4, 1.8.2.3, nor 1.8.5.8, parts of the sulfide oxidation pathways, nor E.C. 1.16.9.1, part of Fe(II) oxidation pathway despite the positive correlation with these metabolisms (Figure 8).

Sulfide oxidation was shown as occurring in all sites (Figure 7B), and PICRUST2 results (Supplemental Table 8) match this expectation, although the predicted members performing this metabolism varied. Either sulfide oxidation enzymes bacterial sulfide:quinone reductase (E.C. 1.8.5.4) or hydrogen-sulfide:flavocytochrome *c* oxidoreductase (E.C. 1.8.2.3), responsible for sulfide oxidation (Figure 7A reaction 1), were predicted in 345 ASVs. However, none of these ASVs were found in all sites. Few ASVs with these sulfide oxidation enzymes were present in multiple sites and these were identified as genera *Methylotenera* (9 sites), *Nitrosarchaeum* (7 sites), or various ASVs unidentified below the family level of Comamondaceae (Supplemental Table 8). Thiosulfate oxidation or disproportionation (Figure 7A reactions 3, 8, and 9) was predicted in 595 ASVs based on E.C. values for 1.8.2.2, 1.8.5.2, 1.8.5.5, or 2.8.1.1. These included ASVs *Bacteroides, NS11-12_marine_group, Luteolibacter, Polynucleobacter,* and *GKS98_freshwater_group* which presented a positive correlation to SO (Figure 8).

Members present in some DSR metabolism tests with PICRUST2 results for adenylyl-sulfate reductase (Supplemental Table 8 EC 1.8.99.2) included *Gallionella, Desulfurivibrio, Limnobacter, Desulfosporosinus, Sulfurifustis,* and *Fusibacter.* None of these were present in 3 or more samples to have Spearman correlations analyses in Figure 8, suggesting site specific reactions. Enzyme sulfate adenyltransferase (Supplemental Table 8 EC 2.7.7.4) was present in 770 of the Riverton ASVs but was not considered for verification against metabolism test due to its conflicting presence in various assimilatory sulfate reduction pathways.

Specific members are shown as correlated with metabolisms of interest, although the strength of most correlations in Figure 8 is lower than geochemical-ASV correlations in Figure 4B. This paired with PICRUST2 results show sites as having most general metabolic pathways shared, and specific ASVs are predicted as performing parts of pathways of interest.

## 4 Discussion

### 4.1 Microbial community membership varies across the Riverton site based on location, compartment, and geochemistry

The Riverton, WY former uranium processing site has been geochemically^64,65^ and hydrologically^8,66^ characterized, results of which have driven site modeling^67^ and remediation plans^19^. There were a few studies that investigated specific microbial reactions of interest^68^ or assessed near-river region SSMA microbial community by depth^23^, however, microbial community membership and reactions have not been assessed throughout the site.

In measuring and comparing microbial membership across the former tailings area (FTA), near-river region (SSMA), upgradient, cross-gradient, and surface, the results validated the first hypothesis that the microbial community varies across the Riverton site. There were only 27 ASVs that fulfilled the requirement of presence in 3 sites when filtering samples for inclusion in statistical analyses that generated heatmaps (Figure 4). The dispersed beta diversity (Figure 2) further indicated heterogeneity of the microbial community. This is also supported by the striking differences at various taxonomic levels between surface waters and groundwater, between regions of the aquifer, and between sites, as visualized in the taxa bar plots (Figure 3) and discussed in section 3.2.

The planktonic community was identified as having different abundances from soil/sediment-bound community results previously published. Riverton SSMA region soil microbial community profile from the topsoil through the aquifer identified Proteobacteria as comprising 43 – 50% of the total community when including the class Deltaproteobacteria, which was the predominant class of Proteobacteria found by that study in the aquifer^69^. Since that article has been published, the Deltaproteobacteria class has been split from the Proteobacteria phylum and four new phylum are now considered: Desulfobacterota, Myxococcota, Bdellovibrionota, and Desulfobacterota^70^. Because of this rearrangement, the comparison of Proteobacteria levels and abundances found in this study are not as direct. The relative percentage of planktonic Proteobacteria measured in an SSMA region near the location of the previous study soil profile had similar abundance, ranging from 48 – 64% (Figure 3), despite Deltaproteobacteria now being considered a separate phylum. Phylum level differences between the aqueous planktonic (this study) and soil (previous study) communities are also notable for Bacteroidetes, which ranged from 4 - 8% in SSMA region soil^23^ whereas in the SSMA groundwater values were much smaller, between 0.2 – 1.8% (Figure 3, Supplemental Table 5). Although the number of studies and samples are limited, this suggests that in addition to location differences in microbial community driven by geochemistry, the microbial community within one location have different membership in the planktonic aqueous phase and the attached phase within soils or sediments, consistent with other study results^25^.

Both CCA and Spearman correlations between geochemistry and shared ASV abundance show significant multiple positive associations between alkalinity, conductivity, sulfate, Mo, U, Mn, and Fe with a negative pH correlation (Figures 4). Another Riverton-site specific literature also reported community variability based on covarying factors of depth, moisture, and metals concentrations at near-river sampling cores^23^. Other uranium contaminated sites also have reported patterns of cross-site microbial heterogeneity and association with geochemical parameters including: Oak Ridge, TN where it has been shown that 16S rRNA data is able to distinguish between U contaminated and uncontaminated sites^26^ and Rifle, CO where it was shown that shorter physical distance differences between microbial communities were ascribed to geochemical or topographical differences between sites^25^.

The Riverton soil community membership has been shown by another study to remain stable over seasonal hydrological and geochemical redox conditions, whilst varying by depth.^23^ Although membership varied, many locations tested in this study displayed capacity for the same metabolisms (Figures 5, 6, & 7), suggesting shared metabolic pathways enabling the community to respond to seasonal changes by altering metabolisms performed, e.g. using different terminal electron acceptors (e.g. oxygen, nitrate, Mn, and Fe).

### 4.2 Metabolism results across the site and relationship to uranium oxidation state

We also hypothesized that the microbial community reduction-oxidation metabolism potential will vary spatially, with activities of and members with known reductive metabolisms being found closer to the naturally reduced zone by the Little Wind River, and more oxidizing metabolisms found in the FTA region. Our results partially support this hypothesis, for although the microbial community varied by geochemistry (Figure 4), metabolism results generally varied less. It is possible that these microbial members that perform metabolisms with minimal results did not survive pumping, transport, or enrichment test preparations e.g. from oxygen toxicity to some S-reducing microorganisms^71^ impacting their ability to grow in the laboratory medias. Furthermore, the well-known “Great Plate Count Anomaly”^72^ of minimal environmental microorganisms being able to grown in laboratory conditions will lead to only specific members to grow outside of their natural environment and display phenotypes of interest. Others have shown that S- and Fe- reducers have higher composition in sediment-associated communities than the planktonic^24^. Therefore, microorganisms with phenotypes of interest not found in the planktonic community were present in the sediment-bound community metagenome. Overall, the Riverton aqueous microbial community displayed equal potential across the site for NR (Figure 5C), MnR (Figure 6C), and SO (Figure 7B) metabolisms, and similar levels for FeR metabolism (Figure 6A). Convincing evidence of direct microbial enzymatic reduction of U(VI) metabolism and AO metabolism was not seen in any sites (Figures 6E and 5B).

Riverton sites with a similar set of detected microbial reduction-oxidation metabolisms in this study were grouped, and these reactions along with their relationship with U oxidation or reduction were visualized in Figure 9. Three surface water and three groundwater groups emerged from variabilities in FeR, FeO, and DSR. FTA groundwater sites 784 and 859-3 nearest the retention pond were grouped based on FeR, DSR, and FeO capability. Groundwater cross-gradient site 720, FTA site 860-3, and all SSMA sites were grouped based on FeR capability and inability of DSR and FeO metabolisms. This does not support the hypothesis of more reductive metabolisms occurring nearer to the naturally reduced zone by the Little Wind River, for although the oxbow lake near the river demonstrated DSR, the SSMA groundwater sites did not. Rather, the groundwater sites in the FTA nearest the retention pond demonstrated this metabolism. The hypothesis that oxidative metabolisms being found in the FTA is mostly supported. SO was found throughout the aquifer and FeO was found at FTA sites 784 and 859-3, however AO was not found at any site of the aquifer.

**Figure 9.**
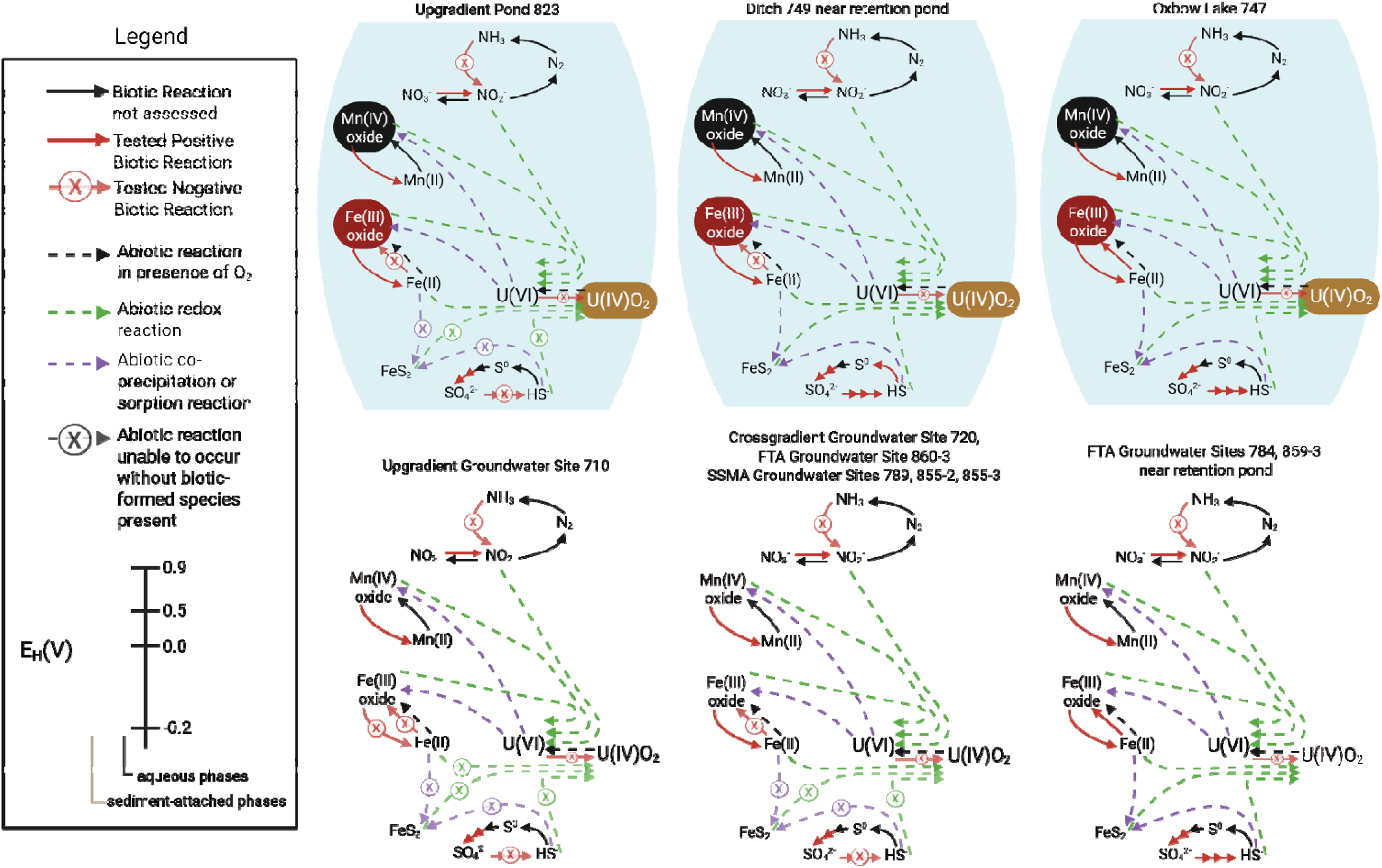
Microbial community metabolic pathway and potential for uranium interaction models of Riverton site regions. Surface water sites are depicted in blue with suspended particles as filled ovals. Groundwater sites are demonstrated as pore-spaces, with sediment-associated phases in brown-regions and generally aqueous phases in the center of each. Biotic reactions are solid lines and abiotic reactions dashed lines.

Nitrate reduction metabolism is of interest because nitrate is a frequent co-contaminant with U,^73^ and microbial nitrate reduction to nitrite (Figure 5A, reactions 1 and 4) can indirectly oxidize and mobilize U(IV) (Figure 9)^37^. Depletion of nitrate was seen in all results (Figure 5C), and both nitrate reduction I (DENITRIFICATION-PWY) and VII (PWY-6748) were predicted in all sites from ASV results by PICRUST2 (Supplemental Table 9). One or more EC genes part of these pathways were present in 364 ASVs, however no ASV was predicted with all denitrification genes (Supplemental Table 8). This supports the idea of microbial communities functioning collectively to support processes like denitrification. Cross-site potential for nitrate reduction matched results from other sites; nitrogen metabolisms in groundwater communities have high redundancy although the major members present vary^74,75^. Denitrification metabolism evaluation at Oak Ridge, Tennessee also found that this metabolism was stimulated with addition of a carbon source^36^. Another study using sediments from Shiprock, New Mexico showed microorganisms capable of multiple anaerobic metabolisms preferentially reducing nitrate prior to Fe- and U- reduction, and that nitrate can act as an inhibitor for these metal reductions^76^. At the Riverton site, nitrogen as nitrate and nitrite is at relatively low concentrations in the groundwater; the last reported measurements for all groundwater values were between 0.01 – 2.5 mg/L and most values within the U plume were less than 0.4 mg/L (2015 measurement)^28^. This is far below other DOE managed sites where nitrate and nitrite as nitrogen have been more recently measured, e.g. Rifle CO Disposal site with levels up to 26.2 mg/L (2024 measurement)^28^ and Tuba City AZ Disposal site with levels up to 725 mg/L (2025 measurement)^28^. Therefore, although nitrate reduction capacity was ubiquitous in surface and groundwaters in Riverton, nitrate reduction as a source of nitrite to abiotically oxidize reduced uranium is likely not an important biogeochemical pathway at this site.

Another potential source of uranium-interacting nitrite is ammonium oxidation which produces nitrite as an intermediate (Figure 5A, reaction 5) prior to oxidation of nitrite to nitrate. Despite that other studies have found ammonium oxidation present in the Riverton aquifer microbial genome above, at, and below the water table in sediment^77^ and similar environments based on *amoA* gene abundance^68^, metabolism results did not identify AO as occurring (Figure 2B). Predicted pathway coverage identified two ASVs, both genera *Nitrosomonas,* with both aerobic ammonium oxidation I enzymes (Supplemental Table 8, EC 1.14.99.39 and 1.7.2.6). These ASVs were present in only sites 747 and 784. 16 other ASVs were predicted to have either enzyme in the genome, but not both. This limited predicted ability to perform ammonium oxidation does not match previous metagenomics results. The previous study^77^ tested sediment-bound microbial communities, so this may indicate a difference between planktonic and sediment-bound microbial communities, 16S rRNA amplicon sequencing based predictive behavior versus metagenomic analysis, or a difference between microbial community potential genomic ability and activity displayed under conditions tested. Due to the lack of evidence of directly measured AO metabolism in the present study, the microbial models in Figure 9 are not shown with AO metabolism.

Biogenic and chemically-precipitated manganese oxides can oxidize reduced U(IV) and bind U(VI)^78^. Mn reduction assay showed complete depletion of Mn(IV) in all sites (Figure 6C and 9) despite no prediction of the PICRUST-identified manganese reduction pathway (Supplemental Table 8, EC 1.16.2) in the Riverton community ASVs. Furthermore, aqueous Mn concentrations were shown in Table 2 and Mn(II) species were predicted with PHREEQC (Supplemental Table 4) across all sites tested, suggesting that active Mn reduction occurs at the site. A limited capacity for Mn oxidation was predicted, with 161 ASVs predicted capable of Mn oxidation II (Supplemental Table 8, EC 1.16.3.3). Mn oxidation tends to occur biotically as it is thermodynamically favorable in oxidizing conditions, but requires a high activation energy and is thereby often kinetically controlled by microorganisms^22^. Overall, Mn was shown as mostly reduced and microbial communities as contributors to Mn(II) generation.

Our results suggest that microbial iron cycling metabolisms occur at the Riverton site and previous literature detail many routes of Fe interaction with U. Fe reduction occurred at all sites except upgradient site 710 (Figure 9), which is outside of the groundwater plume. Aqueous Fe was detected at all plume locations (Table 2), suggesting that Fe reduction is actively occurring at the site. Biotically reduced Fe(II) can contribute to partial abiotic reduction of U(VI) to U(IV)^79^, but this typically occurs once the aquifer has reached sulfate-reducing conditions where biogenic FeS species act upon U(VI)^80,81^. Furthermore, as the U species are mostly Ca-uranyl-carbonates (Supplemental Table 4) which have lower redox potentials than Fe(III)^18^ and decreased reduction in the presence of ferrihydrite^82^, it is unlikely for U(VI) reduction to occur prior to Fe(III) reduction. Fe(III) oxides can also contribute to uranium oxidation^83^. At Riverton, U has been shown to associate with metal oxides including Fe and Al in the former tailings area^64^. This matches with other sites which have reports of U(VI) association with iron oxides including ferrihydrite^84^, goethite, and hematite^17^. If the microbial community is stimulated to reduce Fe, this could lead to an initial increase in aqueous U(VI) from desorption of U from Fe oxides. However, if the aquifer’s reduction potential drops further causing more reducing metabolisms UR or DSR to occur, U will become reduced to uraninite or other less mobile U phases. Three sites, oxbow lake 747 and FTA near-retention pond sites 784 and 859-3, were shown as capable of Fe oxidation (Figures 6B & 9). These were mostly the same sites where biotic dissimilatory sulfate reduction occurred in metabolism tests (Figures 7 & 9), however these reactions are not thermodynamically favorable to be combined/paired^85^ but rather occur in differing conditions. The impacts of sulfur metabolisms on U speciation are discussed next.

Microbial sulfur metabolisms have well established reactions that influence uranium speciation and are of interest at the Riverton site due to elevated sulfate concentrations (Table 2) that correlates with microbial membership (Figure 4, discussed in results section 3.3). Stimulation tests for sulfide oxidation were consistently positive (Figure 7) across the Riverton site. In the context of groundwater, SO is of interest as sulfide oxidizing bacteria can be an important source of organic C in otherwise limited nutrient regions^86^. Recently, it has been shown that SO can be coupled to FeR and occurs at faster rates biotically than the abiotic reaction alone^21^, showing that these two cycles can be linked biogeochemically and suggests that other metal metabolisms may be acting together in ways not previously considered.

The major sulfur reduction pathway in groundwater is dissimilatory sulfate reduction (Figure 7A, reaction 5), and sulfate reduction hotspots in fine-grained sediments contain Fe(II) and S(II) which can act as electron donors to bind and reduce U, causing U(IV) accumulation^80,81^ (Figure 9). Perzan’s 2021 Riverton site modeling^67^ showed S reduction being key for redox condition establishment and maintenance alongside fluid residence time as controlled by aquifer permeability. As S reduction results varied across the microbial community, this will vary if and how U reduction occurs. Sulfate reduction was detected in the oxbow lake, ditch near the retention pond, and FTA sites 784 and 859-3 nearest to the retention pond (Figures 7 and 9). During previous sampling in May 2023, measured groundwater oxidation-reduction potential (ORP) values were lowest for the near-retention pond region of the FTA (-241.9 – -125) compared to higher values nearer 860-3 (-38.3 – +10.1) (internal DOE Legacy Management database). This matches S reduction capability of the microbial community; generally S reduction occurs at lower ORP levels where other electron acceptor abundances are depleted^87^. 81 ASVs were predicted to have dissimilatory sulfate reduction ability based on presence of adenylyl-sulfate reductase (Supplemental Table 8, EC 1.8.99.2). These ASVs varied, and none were present in all sites with noted sulfate reduction nor multiple sites as discussed in section 3.6, suggesting that different members were responsible for the observed DSR metabolism.

### 4.3 Microbial Reactions and Relationship with Remediation Efforts

Typical remediation strategies for U remediation rely on bioreduction, where U is reduced either enzymatically by the microbial community or indirectly by biogenic reduced Fe- and S products through C inputs^14–16^. The community at the Riverton site had mixed ability to completely reduce U and S (Figures 6, 7, & 9). Furthermore, the aquifer is susceptible to re-oxidation through flooding-drying events^6^. Therefore, it is unlikely that bioremediation relying on U immobilization would work as intended for long-term stabilization of reduced U.

Oxidative enhanced flushing has been trialed at the site to increase U flushing from the aquifer. Oxidative enhanced flushing would most likely have the highest impact on areas that demonstrated microbial sulfate reduction and therefore U reduction, e.g. FTA sites nearest the retention pond (Figure 1). During enhanced oxidative flushing events to the saturated zone of the SSMA, changes to U concentration appeared to follow expected changes based on tracer concentrations of dilution alone^19^. As the SSMA planktonic community was not shown capable of direct U reduction (Figure 6) nor DSR metabolism (Figure 7) which could result in reduced U (Figure 9), this suggests that U is likely found in an oxidized state. This is further supported by geochemical modeling (Supplemental Table 4) showing mainly U(VI) species across the site. This would explain the SSMA result as oxidation would not occur to already oxidized U species. However, the FTA saturated zone injections did appear to have an oxidation event occur as U levels rose above initial condition levels^19^. Interestingly, although FTA sites nearest the retention pond showed feasible DSR metabolism, FTA community 860-3 closest to the location of the injection experiment did not show this metabolism. Sultana et al., 2024^64^ found U associated with Fe-rich coatings at this site, suggesting that U sorption to Fe as an important factor in this region. Aqueous Fe(II) presence (Table 2) and Fe reduction by microbial communities (Figure 6) were observed, so microbial Fe activity if changed would influence associated U conditions. Further studies will be required to understand the mobilization factors responsible for this uranium increase during oxidative enhanced flushing.

## Conclusion

This study of the aqueous phase microbial community membership and stimulable metabolisms across the Riverton, WY former uranium processing site confirmed the hypothesis that the microbial community varies by site geochemistry, especially sulfate, conductance, and pH acting as primary drivers. Contrastingly, most of the microbial communities displayed capabilities to utilize nearly the same set of transition metals and inorganic molecules as terminal electron acceptors (O_2_, NO_3_^2-^, Mn(IV), Fe(III)), despite having almost completely distinct microorganisms present. This highlights the resilience of Riverton microbial communities in that they are primed to handle the changing redox conditions in this suboxic and variably reduced aquifer. More broadly, it is another observation of how sequencing alone would not account for system phenotypes and why measuring microbial community activity in conditions replicating *in situ* conditions is crucial for making management and treatment decisions. Moreover, the instances of uncommonly displayed metabolic capacity in our experiments (Fe oxidation, Fe reduction, and S reduction) contradicted our second hypothesis of reducing metabolisms being found primarily in the near-river reduced zones at Riverton. Although not connected through a biogeochemical cycle, the highest capacity for Fe oxidation and S reduction were found at the same site but not in an expected location based on site redox. This again points to the ability of the community to respond to fluctuating conditions. When we combined our microbial results with the site geochemistry and U speciation, we were able to provide an explanation for the lack of U enhanced oxidation during oxic injection observations in the SSMA, as S and U reduction were not found at this site. In the FTA at the site of oxic injections, we found that Fe(III) reduction is a key component of U transport as microbial Fe(III) reduction was observed and U has previously been characterized as associated with Fe. As Fe is reduced by microbial activity to a more aqueous form, this could free associated U where S reduction is not observed. The results of this study provided microbial insights into and potential explanations for the documented efficacy of applied uranium remediation strategies at the different locations within the Riverton site, showing that a uniform cross-site management approach will likely not be effective. Although this study is site-specific, the findings are applicable to other variably reduced, metal-contaminated aquifers.

## Author Contributions

*C.J.Par. refers to author Charles J. Paradis and C.J.Pet. refers to author Catherine J. Pettinger* Conceptualization, C.J.Par. and E.L.-W.M.; Data curation, C.J.Pet.; Formal analysis, C.J.Pet.; Funding acquisition, R.H.J., C.J.Par. and E.L.-W.M.; Investigation, C.J.Pet. and A.M.W.; Methodology, C.J.Pet., A.M.W., R.H.J. and E.L.-W.M.; Project administration, C.J.Par. and E.L.-W.M.; Resources, R.H.J., C.J.Par. and E.L.-W.M.; Software, C.J.Pet.; Supervision, R.H.J. and E.L.-W.M.; Validation, C.J.Pet.; Visualization, C.J.Pet.; Writing – original draft, C.J.Pet. and E.L.-W.M.; Writing – review & editing, C.J.Pet., A.M.W., R.H.J., C.J.Par. and E.L.-W.M.

## Data Availability Statement

All data are found in the manuscript and Supplementary Materials except raw sequencing files which have been uploaded to NCBI as BioProject PRJNA1365257 and are available at the link: https://www.ncbi.nlm.nih.gov/bioproject/PRJNA1365257.

## Acknowledgments

This project was funded by the National Science Foundation Chemical, Bioengineering, Environmental and Transport Systems Award 2229870, Enhanced Biogeochemical Flushing of Uranium in Groundwater.

The authors thank RSI Sampling Team including Sam Campbell for sampling, field measurements, and shipping of samples. The authors gratefully acknowledge the assistance of James Lazarcik and Sarah Hanes for use of facilities and instrumentation in the UW-Madison Core Facility for Advanced Water Analysis. The authors recognize support from the Ricke Lab Sequencing Center within the UW-Madison Department of Animal and Dairy Sciences for 16S amplicon PCR preparation and sequencing. The authors also thank the Rey and Ricke labs for anaerobic chamber usage. The authors thank Dr. Patricia Tran for bioinformatics support on the UW-Madison Center for High Throughput Computing. The authors thank Drs. Ginder-Vogel and Roden for feedback and discussion on results. Graphical abstract and Figure 9 were created in https://BioRender.com.

