## Supplemental Figures for "Aquifer microbial communities differentially display metabolisms capable of secondary effects on uranium speciation across a former metal processing site"

**Supplemental Figure 1.** Metabolism Tests at 6 week timepoint. In each image, the left tube is the media control. Center-left tube is replicate-1 of metabolism test for the labelled site. Center-right tube is replicate-2 of metabolism test for the labelled site. Right tube is the site-specific sterile control for the metabolism test.

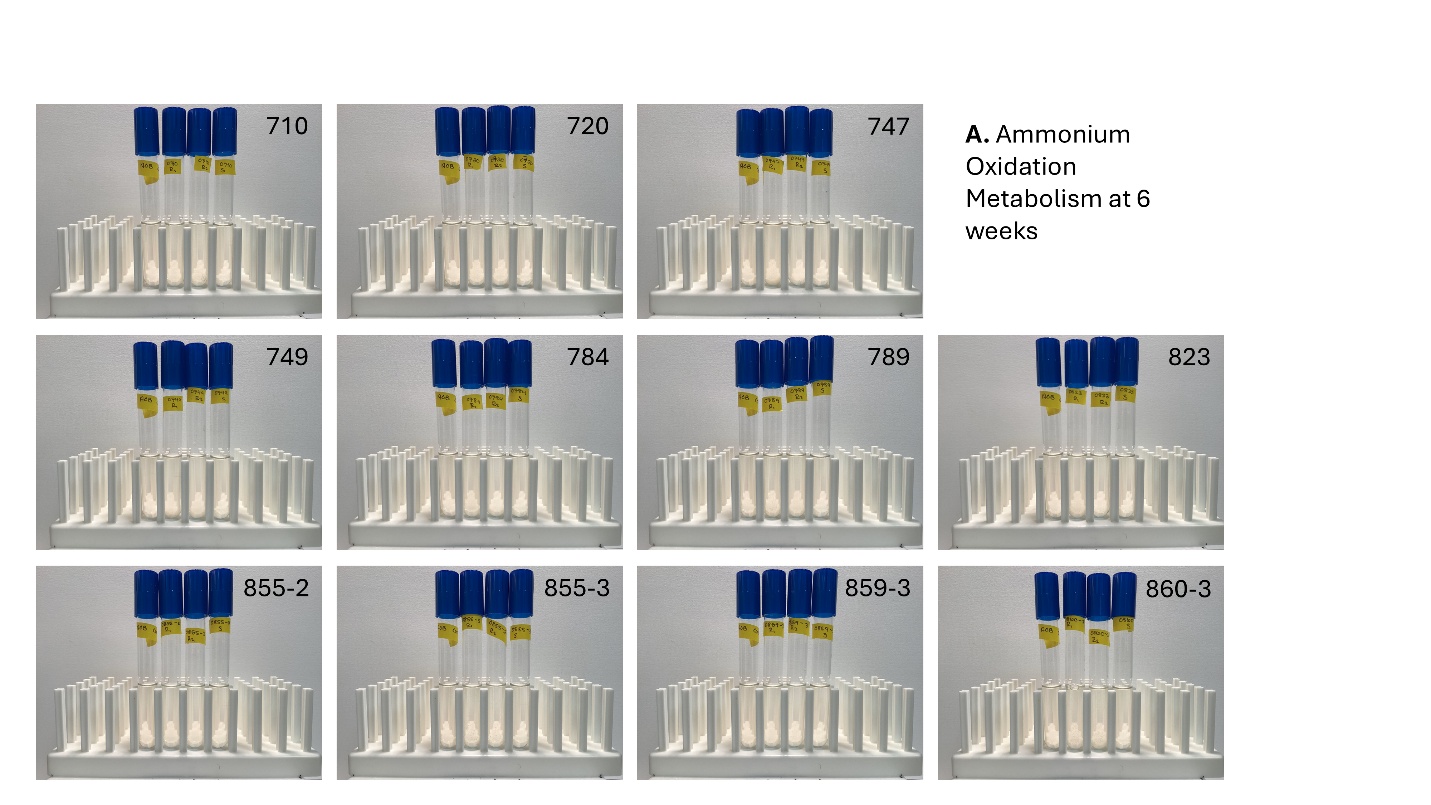

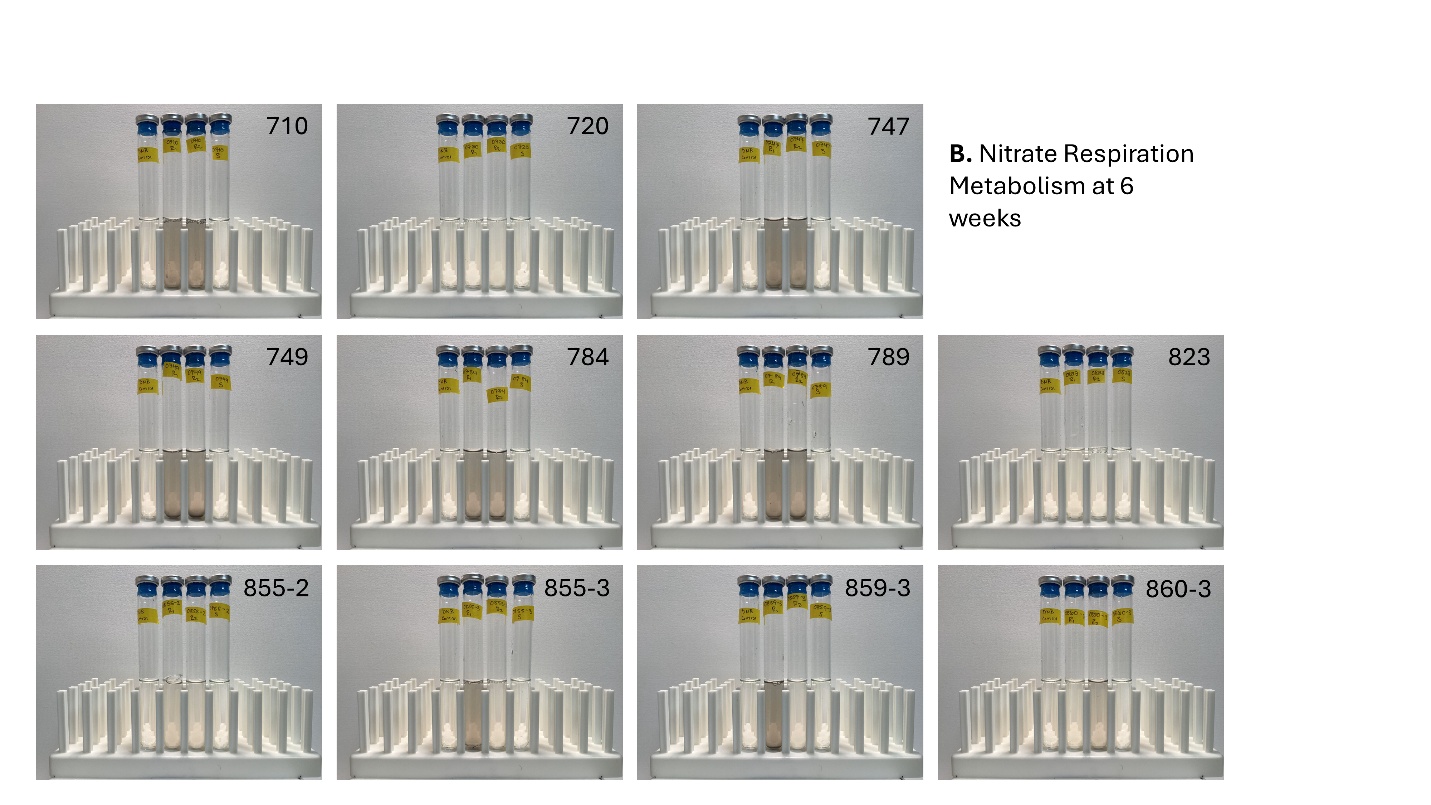

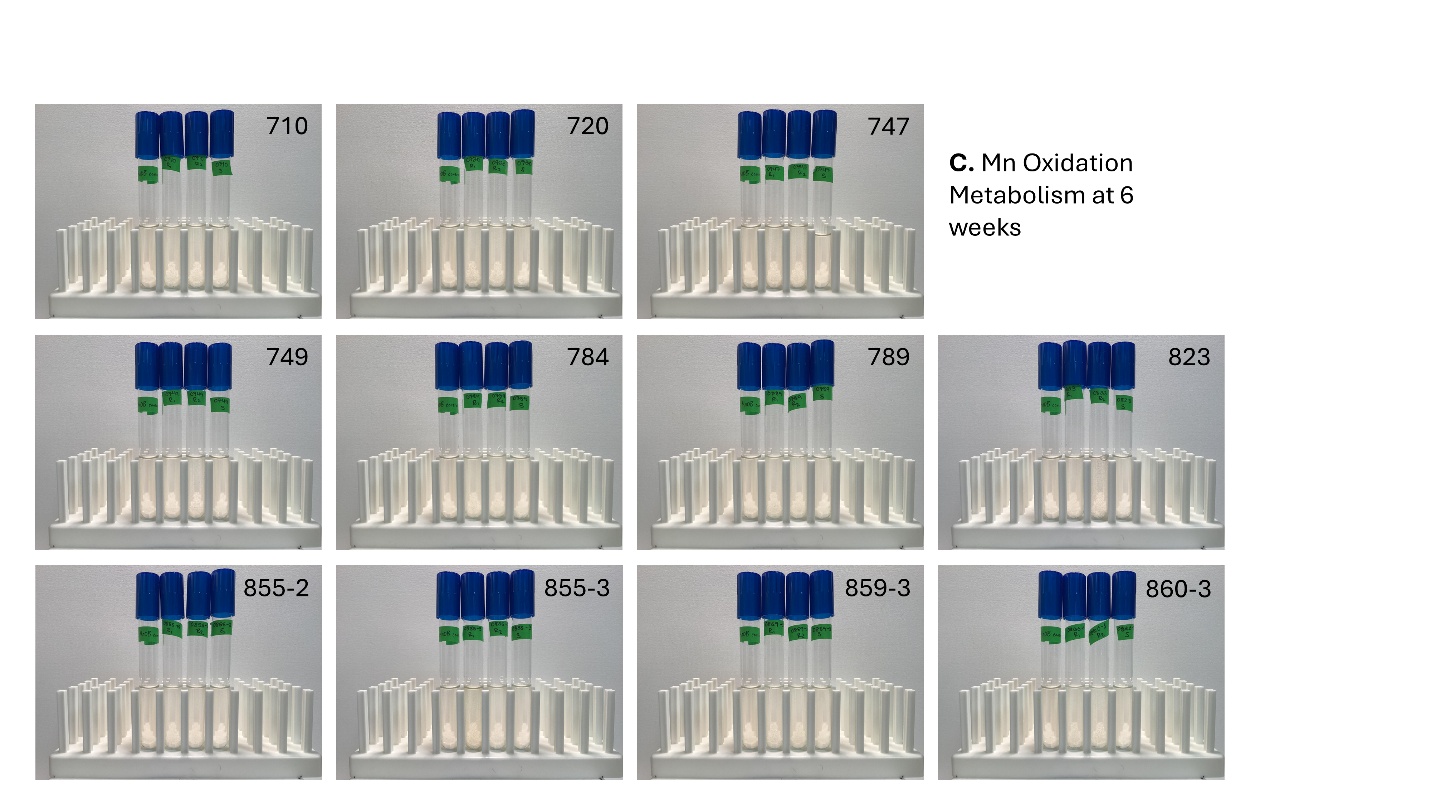

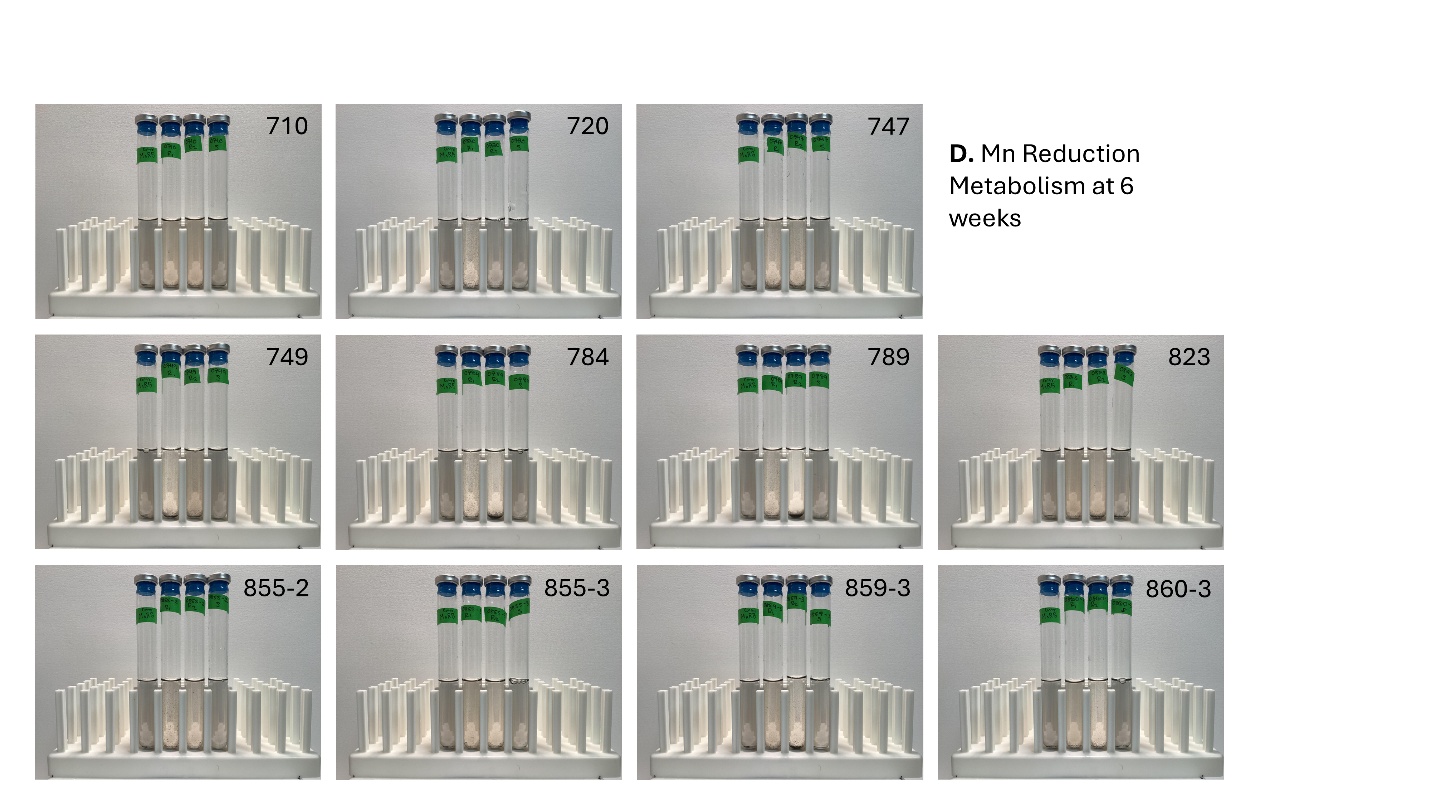

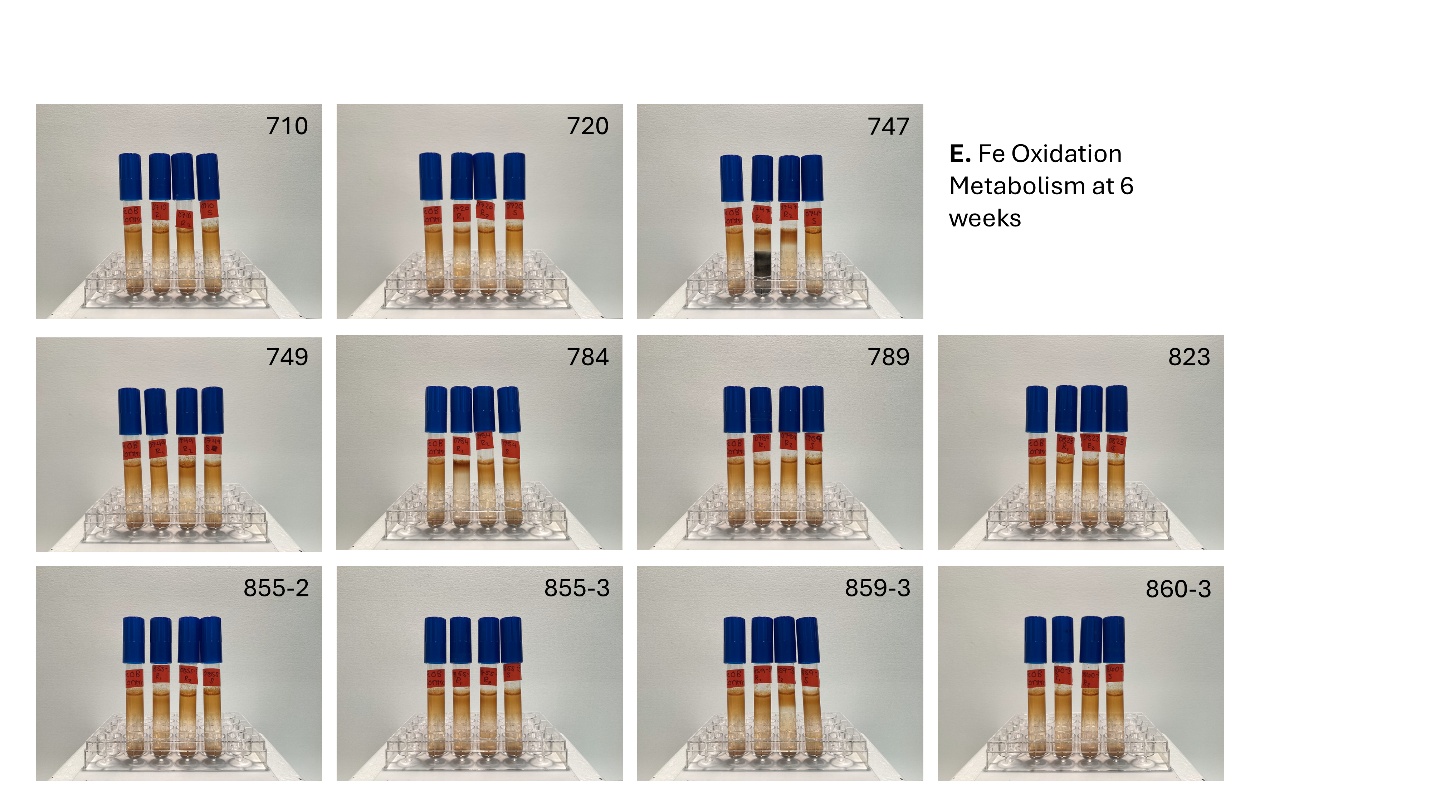

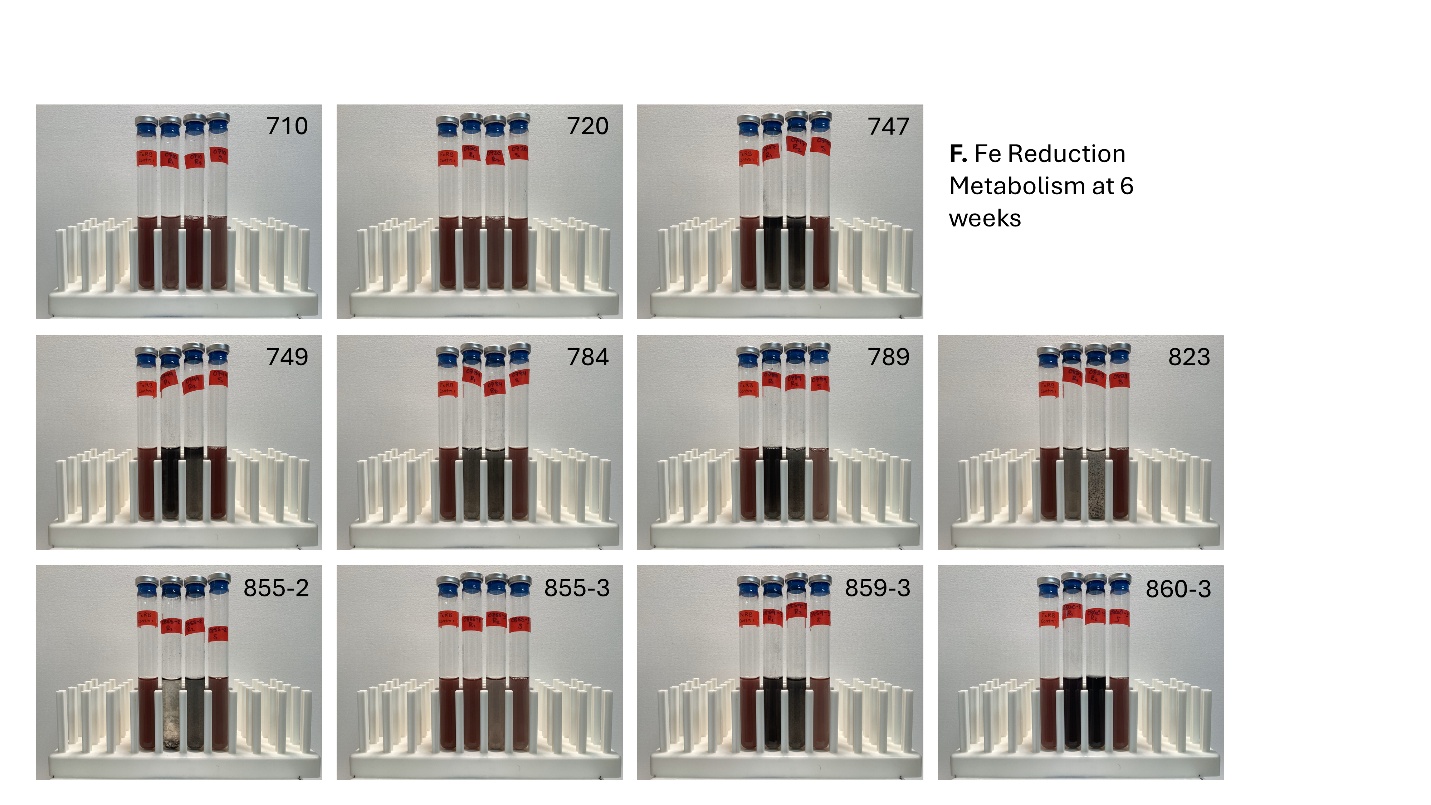

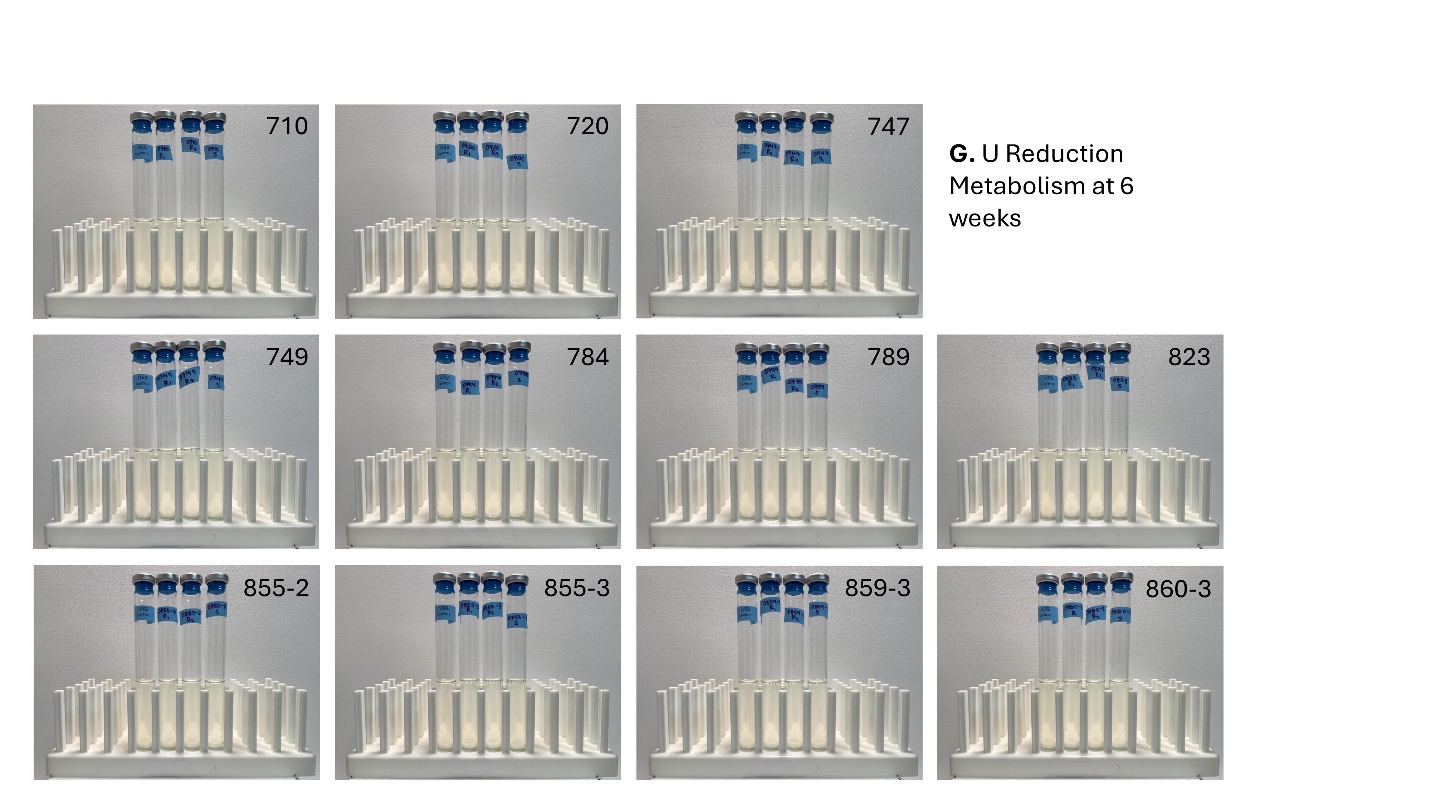

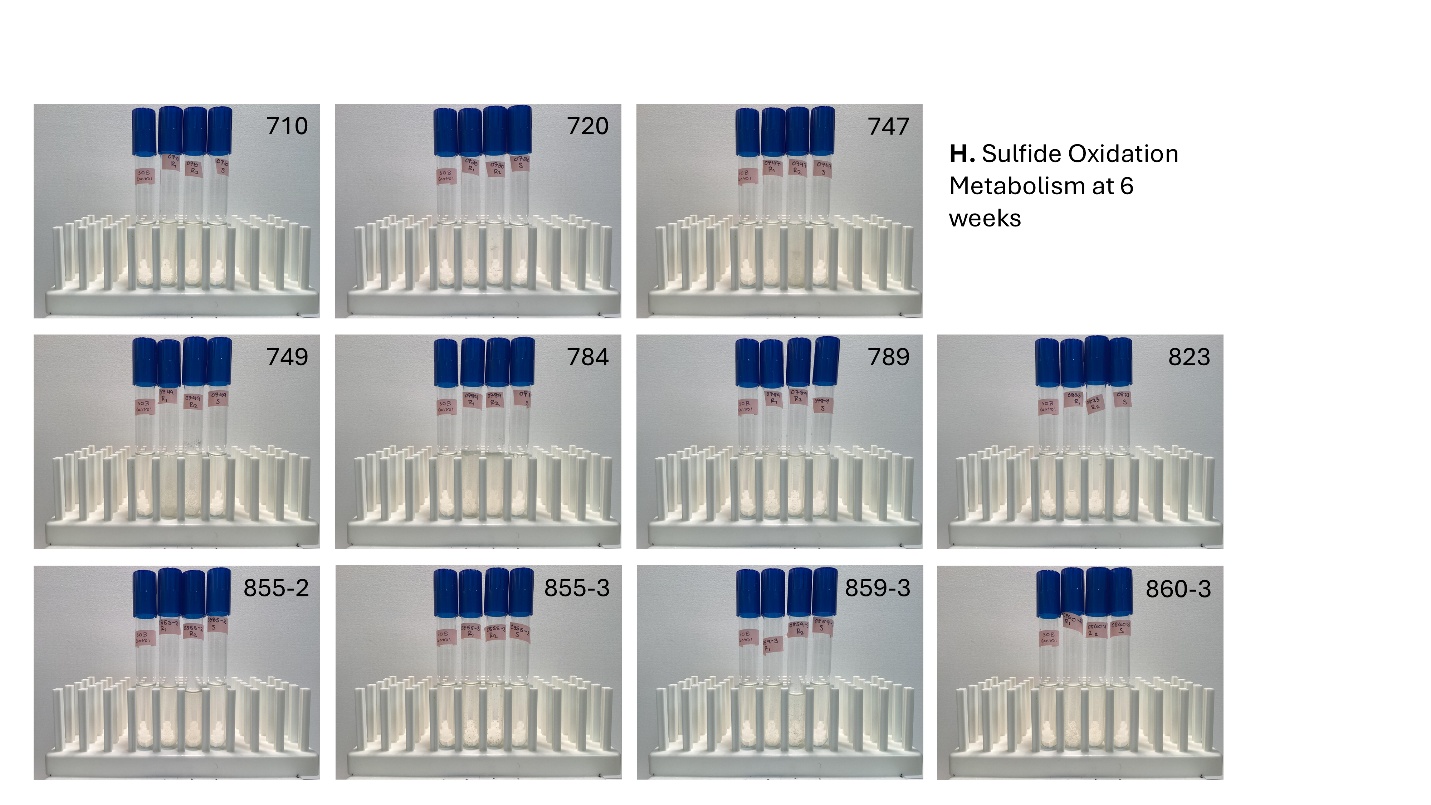

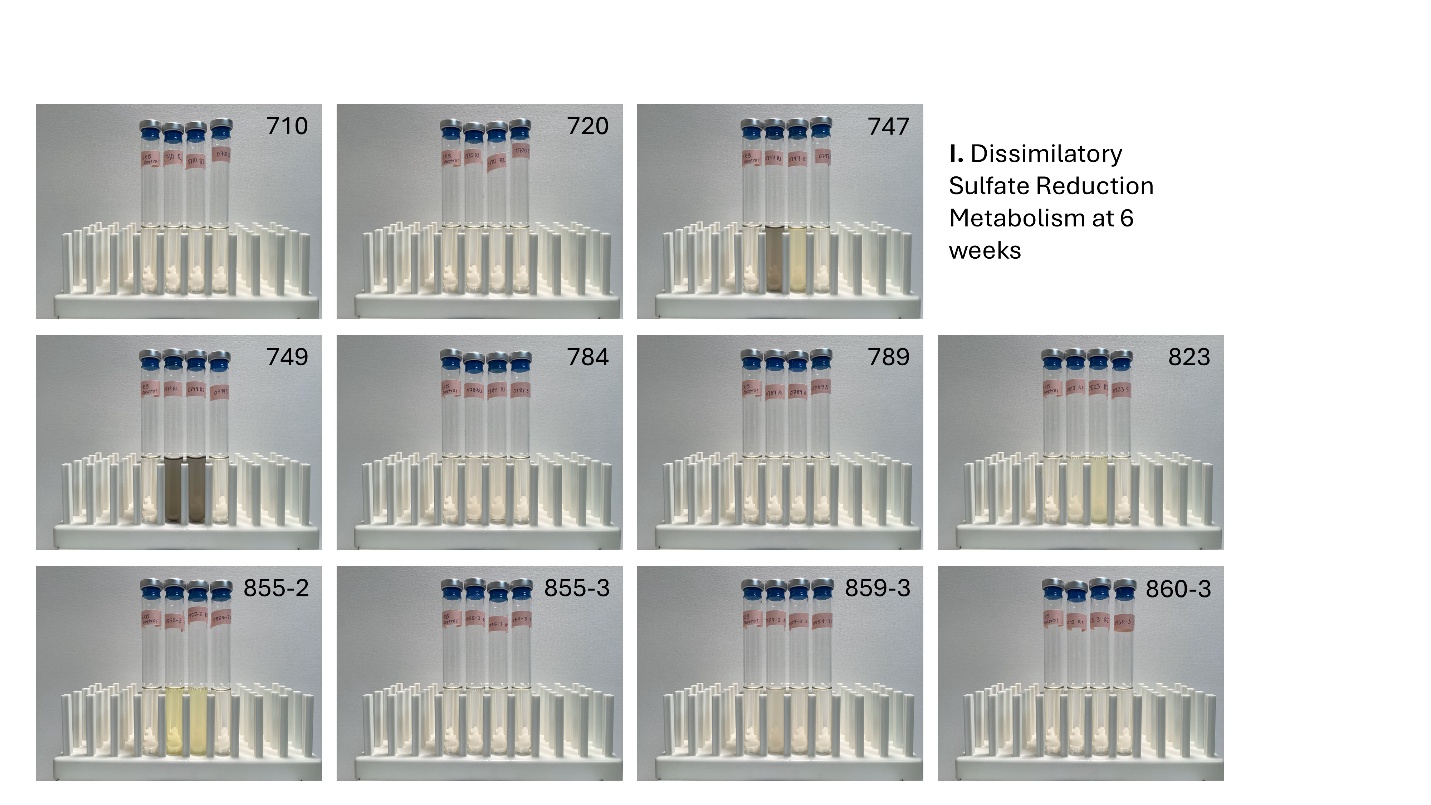
